# Solid-state enzymatic hydrolysis of mixed PET-cotton textiles

**DOI:** 10.1101/2022.07.29.502078

**Authors:** Sandra Kaabel, Jane Arciszewski, Tristan H. Borchers, J.P. Daniel Therien, Tomislav Friščić, Karine Auclair

## Abstract

Waste polyester textiles trap copious amounts of useful polymers, which are not recycled due to separation challenges and partial structural degradation during use and thermo-mechanical recycling. Chemical recycling of polyethylene terephthalate (PET) through depolymerization can provide a feedstock of recycled monomers to make “as-new” polymers, and reduce the accumulation of plastic waste in landfills. Enzymes are highly specific, renewable, environmentally benign catalysts, with hydrolases available that are active on common PET textile fibers and on cotton. The enzymatic PET recycling methods in development, however, have thus far been limited to clean, high-quality PET feedstocks, and most such processes require an energy-intensive melt-amorphization step ahead of enzymatic depolymerization. Here we report that high-crystallinity PET in mixed PET/cotton textiles can be directly and selectively depolymerized to terephthalic acid (TPA) by using a commercial cutinase from *Humicola insolens* under moist-solid reaction conditions, affording up to 30 ± 2% yield of TPA. The process is readily combined with cotton depolymerisation through simultaneous application of cellulase enzymes (CTec2^®^), providing up to 83 ± 4% yield of glucose without any negative influence on the TPA yield. The herein presented selective and/or simultaneous enzymatic hydrolysis of PET/cotton textiles in solid reaction mixtures can expand the biocatalytic recycling processes of PET to less-valuable waste materials, and significantly increase its profitability through operating at very high solid-loading (40%), without the need for melt-amorphization.

## Introduction

Out of the 109 million tons of fiber produced in 2020, including all natural (e.g. cotton, wool) and artificially-made fibers (e.g. polyester, polyamide), only 0.5% was sourced from recycled textiles.^1^ Polyester makes up 52% of the global fiber market, and from the 57 million tons of polyethylene terephthalate (PET) fibers produced yearly, 85% is produced directly from fossil-based sources and 15% from mechanical recycling of plastic bottles.^1^ The post-consumer bottle PET polymer (M_w_ >20,000 g mol^−1^) undergoes thermal and hydrolytic degradation during mechanical recycling,^2^ and thus the resulting textiles (M_w_ ≈ 15–20 000 g mol^−1^) are generally not recycled, but are instead discarded at landfills or incinerated.^3,4^ Due to the increasing demand for reclaimed PET bottles^5^ and the environmental burden of discarded garments,^6^ methods for recycling lower grade waste PET and complex mixed materials, e.g. by textile-to-textile recycling, are of critical interest both from environmental and economic viewpoints.^7^

Considering commonly available recycling processes, it would be preferable to avoid creating mixed-material textiles, since separation of fibers for polymer recovery is difficult at the end-of-life of clothing.^6^ However, the production of mixed-fiber textiles, e.g. PET and cotton, will certainly persist owing to the favorable properties of the resulting garments (color fastness, comfort, functionality, ease of care and softness).^3^ Chemical recycling, *i*.*e*. recovery of PET monomers terephthalic acid (TPA) and ethylene glycol (EG) through depolymerization and their subsequent re-polymerization, can provide access to recycled PET polymers of equal quality to that of virgin PET.^8,9^ Several approaches have been developed for chemical depolymerization of PET,^10,11^ including hydrolysis^12,13^ and glycolysis^8,14–17^ of PET fibers, but only a few studies address chemical recycling of PET in mixed-polymer materials.^12,18,19^ Chemical recycling commonly requires reagents, metals, and additives, as well as elevated temperatures and pressures, which makes these methods energy-intensive, environmentally hazardous, and susceptible to unwanted side reactions.

Enzymatic depolymerization can eliminate the need for fiber separation due to the high substrate specificity of these catalysts, and because it offers an opportunity to conduct transformations under mild conditions (*e*.*g*., aqueous reaction media, atmospheric pressure, and temperature up to 65–70°C). For example, selective enzymatic saccharification of cotton from mixed 60/40 cotton/PET materials was shown to yield 50-60% of glucose (98.3% with NaOH/urea pretreated textiles) in 72 hours, which could be converted to ethanol upon subsequent fermentation.^20,21^ Importantly, the purified polyester fibers were structurally unaltered during the process.^20,22^ Hydrolysis of cotton from different types of cotton/PET waste textile was also demonstrated by solid-state fungal fermentation, resulting in purified PET fibers.^23^ The selectivity of enzymatic hydrolysis was further demonstrated by the step-wise recovery of amino acids and glucose from wool/cotton/PET mixed fabrics using proteases and cellulases, and recovering 96% pure PET.^24^

While recent breakthroughs in protein engineering have delivered PET hydrolases with remarkable activity,^25,26^ such enzymes are generally not active on high-crystallinity PET, such as spun PET fibers^27,28^ which are more recalcitrant than, for example, bottle or packaging PET. Melt-amorphization of waste PET by extrusion, currently considered as part of the enzymatic PET recycling processes,^29^ is energy-intensive and restricted to treating monomaterial waste. Previous reports using a cutinase from *Thermobifida cellulosilytica* demonstrated selective hydrolysis of PET from blended plastic packages, achieving 3.1% and 1.3% yields of TPA in three days at 50°C from either PET (10% crystallinity) with 8% polyethylene, or PET (33% crystallinity) with 7.5% polyamide, respectively.^30^

Our group showed that the hydrolysis of high-crystallinity PET (36% crystallinity) by the cutinase from *Humicola insolens* (HiC, Novozym 51032) can be significantly improved by employing the enzyme in moist-solid reaction mixtures with gentle mechanical mixing (mechanoenzymology) instead of standard aqueous solutions.^31^ Therein, common impurities of post-consumer PET (e.g. 10% polypropylene) did not affect the reaction outcome. Periodic mechanical agitation by ball-milling, a process termed reactive aging or RAging,^32–35^ improved the rate of the enzymatic reaction, yielding 20% of TPA in 3 days.

Herein we turn our focus to the enzymatic hydrolysis of pre- and post-consumer PET and PET/cotton textiles, again using mechanically-agitated moist-solid reaction mixtures. We demonstrate that high-crystallinity (up to 46%) PET textiles and mixed PET/cotton textiles can be partially hydrolyzed to TPA (up to 30 ± 2% yield) and glucose (up to 83 ± 4% yield) by the corresponding enzymes, showing the potential for both simultaneous and stepwise depolymerization of PET and cotton under mild, environmentally benign conditions.

## Results and Discussion

### Polyester and polycotton textiles: composition, choice of hydrolase enzymes and reaction conditions

The studied textiles include woven polyester-cotton and polyester pre-consumer and post-consumer textiles. Information on the composition, structure and source of the textiles are presented in Table 1, whereas characterization details are available in the supporting information together with the optical microscopy images (Figures S1, S2). All textiles contain polyethylene terephthalate (PET) as the main polyester type, therefore a cutinase from *Humicola insolens* (HiC, Novozym 51032, sold as a solution) was used as the PET-hydrolase, known to depolymerize high-crystallinity PET cleanly to TPA under moist-solid reaction conditions.^31^ Cotton, present in 0–35%, was hydrolyzed using the Cellic CTec2 cellulases blend, which contains β-glucosidases, cellobiohydrolases, endoglucanases, as well as hemicellulases, and has previously been shown to efficiently convert biomass to fermentable sugars under mechanoenzymatic reaction conditions.^36^ The herein presented mechanoenzymatic hydrolytic reactions proceed in the presence of a minimal amount of water, acting as a reaction lubricant and a substrate (liquid-to-solid ratio *η* = 1.5 μL mg^−1^, which is ca. 15 molar equivalents of water to monomer), affording a very high (40% w/v) solids loading of the reaction mixture. The water, added as a sodium phosphate (NaPi, 100 mM, pH 7.3) buffer solution, absorbs fully to the textile substrates, resulting in moist-solid reaction mixtures in which chemical transformations are enabled through mechanical liquid-assisted grinding (LAG)^37,38^ and static incubation at 55°C (aging). Solid-to-solid transformations, by LAG and/or aging, are free from the solubility limitations typical of bulk solvent media and are often faster than conventional solution reactions due to enhanced opportunity of molecular collisions at high-solids reaction conditions.^39,40^

**Table 1.** Composition and source of the studied textile samples.

| Textile code | Type, composition<sup>[a,b]</sup> | Weight<sup>[c]</sup> | Crystallinity PET<sup>[d]</sup> | Source |
| --- | --- | --- | --- | --- |
| **PRE 65/35** | Pre-consumer<br>65% PET<br>35% cotton | 137.7 g/m<sup>2</sup> | ND | Fabricville (Broadcloth) |
| **PRE 80/20** | Pre-consumer<br>80% PET<br>20% cotton | 217 g/m<sup>2</sup> | ND | Fabricville (Buckram) |
| **PRE 100** | Pre-consumer<br>100% PET | 102.7 g/m<sup>2</sup> | 39% | Fabricville (Lining) |
| **POST 65/35** | Post-consumer<br>65% PET<br>35% cotton | 120.6 g/m<sup>2</sup> | ND | Clothing, beige |
| **POST 100** | Post-consumer<br>100% PET | 184.6 g/m<sup>2</sup> | 46% | Clothing, dark blue |

### PET and cotton are simultaneously hydrolyzed by the HiC and CTec2 enzymes in moist solid mixtures

Mechanoenzymatic reactions were first performed on the mixed 65% PET/35% cotton pre-consumer textile (**PRE 65/35**) to compare the individual and combined activities of HiC and CTec2 (Table S1, entries 1–6). The textiles, cut into 0.7 × 0.7 cm squares, were ball milled in 15 mL stainless steel milling jars in the presence of HiC (0.65% w/w) and/or CTec2 (0.7% w/w) enzymes presence of HiC (0.65% w/w) and/or CTec2 (0.7% w/w) enzymes for 5 or 30 minutes (30 Hz), followed by 7 days of static incubation at 55°C (Figure 1A).

**Figure 1.**
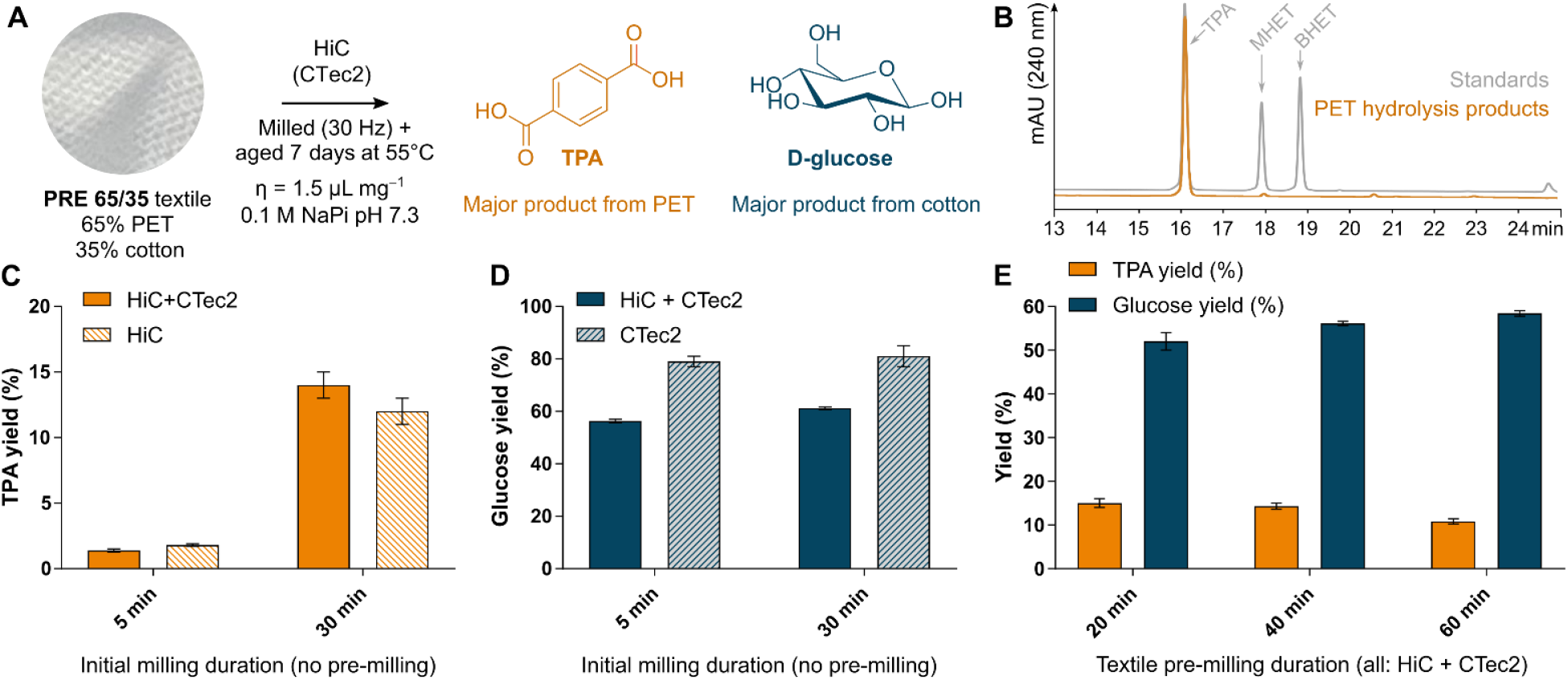
A) General reaction conditions and the major products, terephthalic acid (TPA) and glucose, from the mechanoenzymatic hydrolysis of the PET/cotton mixed textile **PRE 65/35**. Hydrolysis of PET also yields ethylene glycol (yield not determined), and up to 0.5% yield of mono(2-hydroxyethyl) terephthalate (MHET) as the minor reaction product. B) Representative HPLC chromatogram of the PET hydrolysis products (orange), compared to the standards of PET hydrolysis products (grey): TPA, MHET and bis(2-hydroxyethyl) terephthalate (BHET). The minor reaction product (18 min) is MHET, no BHET was detected. C) Influence of applying the HiC enzyme alone (light orange, striped) and in combination with CTec2 cellulases (orange, full color) on the yield of TPA from intact textile, using two different milling durations before aging. D) Influence of applying the CTec2 enzyme alone (light blue, striped) and in combination with HiC (blue, full color) on the yield of glucose from intact textile, using two different initial milling durations before aging. E) Influence of the textile pre-milling duration on the yield of TPA (orange) and glucose (blue) from the mechanoenzymatic hydrolysis of **PRE 65/35** with HiC and CTec2 cellulases simultaneously.

Analysis of the hydrolysis products by HPLC revealed that TPA is the main product of HiC-catalyzed PET hydrolysis, with more than 40-fold selectivity over mono(2-hydroxyethyl) terephthalate (MHET, Figure 1B). The duration of the initial milling step significantly influenced the HiC-catalyzed PET hydrolysis, with a 30-minute milling step resulting in a powdered reaction mixture that reached 12 ± 1% yield of TPA after 7 days of aging, and 14 ± 1% of TPA when co-applied with CTec2, while the more coarse fibrous material from a 5-minute milling step reached only 1.8 ± 0.1% yield of TPA after 7 days of aging, and 1.4 ± 0.1% when co-applied with CTec2 (Figure 1C). Unlike PET breakdown, cotton hydrolysis by the CTec2 cellulases, monitored via glucose assays (see SI for details), was not as significantly affected by the initial milling time (Figure 1D), with aging proceeding after the 5- and 30-minute initial milling to 79 ± 2% (56.4 ± 0.6% when co-applied with HiC) and 81 ± 4% (61.2 ± 0.5% when co-applied with HiC) yield of glucose in 7 days, respectively. It is important to note that the glucose assay represents only the hydrolysis end-product, glucose, after hydrolysis of cellulose by cellobiohydrolases and endoglucanases followed by cellobiose hydrolysis to glucose by β-glucosidase. Therefore, the observed 81 ± 4% yield of glucose indicates an extensive hydrolysis of cotton. We observed that, while the reactions catalyzed by HiC were not significantly affected by the presence of CTec2 cellulases (Figure 1C), the CTec2 enzymes were most efficient in the absence of HiC (Figure 1D).

To gauge the effect of textile pre-micronization on the hydrolysis outcome, dry **PRE 65/35** textile was pre-milled 20, 40 or 60 minutes in stainless steel jars (30 Hz) prior to the start of the reaction. Pre-milling afforded uniform, fine powders (Figure S3), with no intact fibers visible, which allowed excellent mixing of the enzymes and substrates with just a short 5-minute initial milling step, leading to comparable yields of TPA and glucose to the reactions started from intact textiles reacted with 30-minute milling followed by aging, i.e. ca. 15% yield of TPA and ca. 55% yield of glucose (Figure 1E). Compared to longer milling of the reaction mixture, pre-milling the textile has the advantage of decreasing the duration of mechanical action on the enzymes, especially beneficial with enzymes that are less tolerant to milling.

Beige post-consumer textile made of 65% PET and 35% cotton (**POST 65/35**) similarly afforded the highest yield of TPA 8.2 ± 0.2% when the enzymatic reaction was started from a pre-milled powdered textile (Table S2, entry 4), compared to reactions started from the intact material, which gave 1.38 ± 0.09% and 5.8 ± 0.5% yield of TPA for reactions initiated with a 5- and 30-minute milling step, respectively (Table S2, entries 2 and 3). The TPA yield achieved from the **POST 65/35** material was lower in all the tested conditions (Table S2, entries 1–4) than from **PRE 65/35** textile which reached 14–15% of TPA (Table S1, entries 7, 8) and the glucose yield achieved was also lower 41 ± 3% for **POST 65/35** (Table S2, entry 3) compared to 52–56% from **PRE 65/35** at similar conditions (Table S1, entries 7, 8). Similarly, the hydrolysis of a post-consumer 100% PET textile **POST 100**, capped at 8.8 ± 0.2% yield of TPA after 7 days of aging (Table S2, entry 7). The lower yield of TPA may be due to contaminants specific to post-consumer textiles or structural differences inherent to the specific textiles.

### Improved hydrolysis of PET and cotton achieved through Reactive Aging (RAging)

Previous reports on the mechanoenzymatic breakdown of PET^31^ and biopolymers^32–34,36^ have demonstrated that enzyme kinetics under aging conditions exhibit a hyperbolic profile. The initial rates observed during aging, 10.4 mM_TPA_h^−1^ for HiC with PET^31^ and 560 mM_Glc_h^−1^ for CTec2 on corn stover biomass^36^, began to decrease within 48 hours of aging for HiC and within 24 hours for CTec2. With the aim of keeping the reaction rate more stable, we investigated the hydrolysis of textiles with intermittent milling of the reaction mixture (RAging) to improve mixing of the solids during aging. It has been demonstrated that mechanoenzymatic reactions can be accelerated by RAging.^31–34,36^ Based on our previous work with HiC,^31^ the RAging cycle chosen for these enzymatic reactions consisted of 5 minutes of milling and 24 hours of aging at 55°C, repeated for several days (Figure 2A). The yields of glucose and TPA were measured after each cycle.

**Figure 2.**
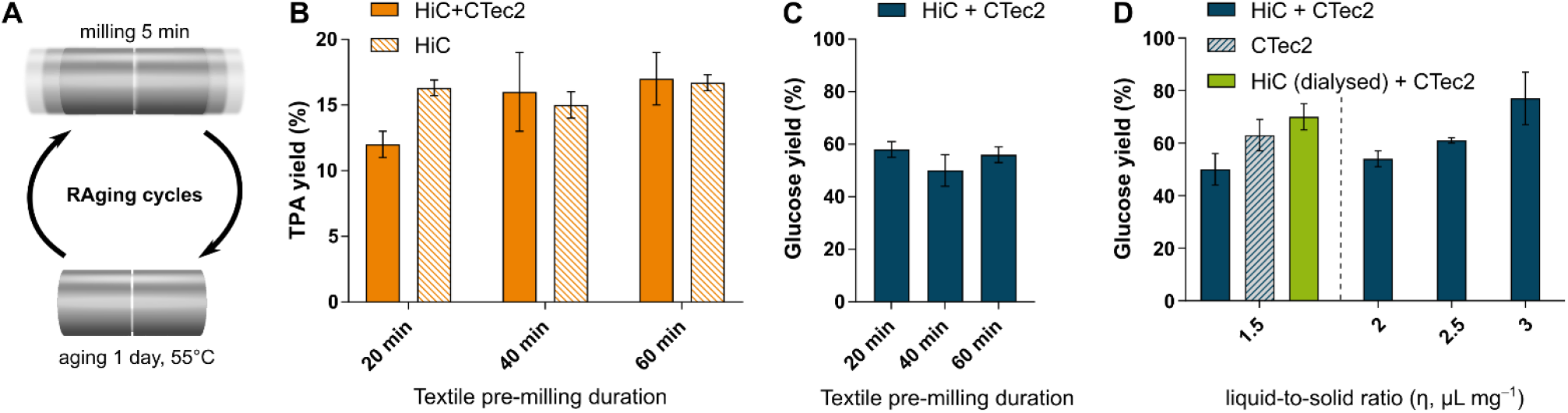
A) RAging regime used, which consists of alternating between periods of ball-milling and static incubation (aging). B) Influence of the textile (**PRE 65/35**) pre-milling duration on the yield of TPA for RAging reactions, showing results for reactions where HiC is applied in the absence (light orange, striped) or in combination with CTec2 (orange, full color). C) Influence of the textile (**PRE 65/35**) pre-milling duration on the yield of glucose for RAging reactions in which both HiC and CTec2 are used (blue, full color). D) Influence of liquid-to-solid ratio (η) on the yield of glucose for RAging reactions in the presence of both HiC and CTec2 (blue, full color), showing also results at η = 1.5 μL mg^−1^ for reactions with CTec2 alone (light blue, striped) or in combination with dialyzed HiC enzyme (green).

While stainless steel milling jars (used so far in this work) efficiently micronized intact textiles, milling of the textiles in poly(tetrafluoroethylene) (PTFE) jars with a single ZrO_2_ ball did not convert fibrous textiles to powder. However, since RAging reactions require the use of PTFE jars, the textiles were pre-milled in stainless steel jars prior to all enzymatic RAging experiments conducted in PTFE jars. Varying the pre-milling duration (20, 40 or 60 minutes) did not significantly influence the outcome of PET hydrolysis from **PRE 65/35** textile under RAging conditions when only the HiC enzyme was used (Figure 2B, Table S3, entries 1, 3 and 6), giving a TPA yield around 15–17%. However, when cellulases were also present in the mixture, the longer 60-minute pre-milling of the substrate yielded 17 ± 2% TPA, while a shorter 20-minute pre-milling led to only 12 ± 1% TPA (Figure 2B, Table S3, entries 2, 4 and 7). In contrast, textile pre-milling for more than 20 minutes did not significantly improve the glucose yield from cotton in RAging reactions (Figure 2C, Table S3, entries 2, 4, 7). Similar to when the reaction mixtures were only milled once, a RAging reaction with CTec2 in the absence of HiC gave a higher yield of glucose (63 ± 6%, Figure 2D, Table S3, entry 5) than when HiC was also present (50 ± 6%, Table S3, entry 4). Interestingly, the negative effect of HiC on CTec2 activity disappears after dialysis, which removes the cryoprotectants and other potential additives present in the commercial HiC solution (see SI for details), affording a 70 ± 5% yield of glucose (Figure 2D). This is consistent with inhibition of the CTec2 cellulases by one or more dissolved components of the commercial HiC enzyme solution, and not by the HiC enzyme itself. Notably, inhibition of cellulases by these dissolved components is also overcome by working at higher liquid-to-solid ratio, which dilutes the commercial HiC solution, and gives a glucose yield of 77 ± 10% at η = 3 μL mg^−1^ compared to 50% at η = 1.5 μL mg^−1^. Lastly, these results also demonstrate the absence of cellulases inhibition by TPA and imply a lack of competition between HiC and the cellulases for the available textile surface.

Previous work has shown that hydrolysis of microcrystalline cellulose and lignocellulosic biomass with *Trichoderma longibrachiatum* cellulases can benefit from hourly milling (instead of daily milling), suggesting that the kinetics of the reaction might be limited by mixing/diffusion of the enzymes in the solid reaction mixture. To test for this, we next investigated the hydrolysis of cotton from **PRE 65/35** textile with the CTec2 cellulases under a RAging regime of 5 minutes of milling and 55 minutes of aging at 55°C repeated over several cycles. Sampling after each cycle demonstrated that such reactions plateaued at a much lower yield of 22 ± 2% glucose in 10 hours (Table S3, entry 12), compared to a similar reaction with daily milling (Table S3, entry 5), which reached 53 ± 2% yield of glucose within the first 24-hour cycle and 71 ± 6% after 3 cycles. This shows that unlike the hydrolysis of biomass substrates,^32^ the reaction rate with textile substrates is not improved by more frequent mechanical agitation of the reaction mixture.

HiC-catalyzed PET hydrolysis of textiles with a smaller cotton content, as in **PRE 80/20** and **PRE 100**, revealed that the TPA yields after RAging also vary (7–17%) depending on the textile (Figure 3A and B, Tables S4 and S5). Surprisingly, the final product of CTec2-catalyzed cotton hydrolysis from **PRE 80/20** proved not to be glucose (Figure 3C, Table S4), therefore the progress of the RAging reactions was also estimated from the weight loss of the textiles (Figure 3D, see SI for details). The highest weight loss was observed for the **PRE 65/35** and **POST 65/35** textiles treated with both HiC and CTec2 (Figure 3D), consistent with the highest yields of TPA and glucose (Figures 3B and C, Table S3 and S6) obtained compared to the other textiles. The measured weight loss of 30–39% for experiments combining the HiC and CTec2 enzymes and 10–13% for experiments using HiC alone match well the expected weight loss calculated based on the yields of TPA, ethylene glycol (assumed 1:1 mol to TPA released) and glucose of the respective reactions (see SI for details). The 15% weight loss of **PRE 100** textile in the presence of both HiC and CTec2 enzymes, also agrees well with the expected weight loss based on the respective TPA yield. The weight loss measured with the **PRE 80/20** textile treated to either HiC only (15.4 ± 0.8%) or to both enzymes (17.6 ± 0.8%) was, however, *ca*. 2.0-fold larger than the expected 8%, based on the corresponding TPA yield and assuming no cotton hydrolysis (Figure 3B, C and D, shown in red). This indicates that, while no glucose was formed from **PRE 80/20**, other soluble hydrolysis products were generated that were extracted in the methanol and water used to collect TPA and glucose, respectively. Subjecting the textiles to the RAging regime without enzymes present did not lead to any weight loss upon washing (Table S7). Furthermore, the limited increase in weight loss observed from **PRE 80/20** when CTec2 was added to HiC compared to when it was not, indicates that the *ca*. 2.0-fold difference between the expected and experimental weight loss is not due to extensive cotton hydrolysis. Indeed, the cotton appears to be largely unreacted in this material, as supported by Attenuated total reflectance Fourier transform infrared (ATR-FTIR) spectra of the remaining solids after hydrolysis of **PRE 80/20** showing no significant differences when HiC is applied alone or together with CTec2 (Figure S9), similar to reactions of the **PRE 100** material (Figure S10). This might be explained by the reduced accessibility of cotton in textiles of higher PET content (PET might not be depolymerized sufficiently to expose the cotton, unless a more efficient PET hydrolases is used). In comparison, the **PRE 65/35** textile shows clear disappearance of the absorbance bands characteristic to cotton (ATR-FTIR: 3337, 2896 and 1160 cm^−1^ for polysaccharide O-H and C-H stretching vibrations and C-O-C asymmetrical stretching vibrations, respectively; see Figure S8) after treatment with HiC and CTec2 (weight loss 38 ± 2%).

**Figure 3.**
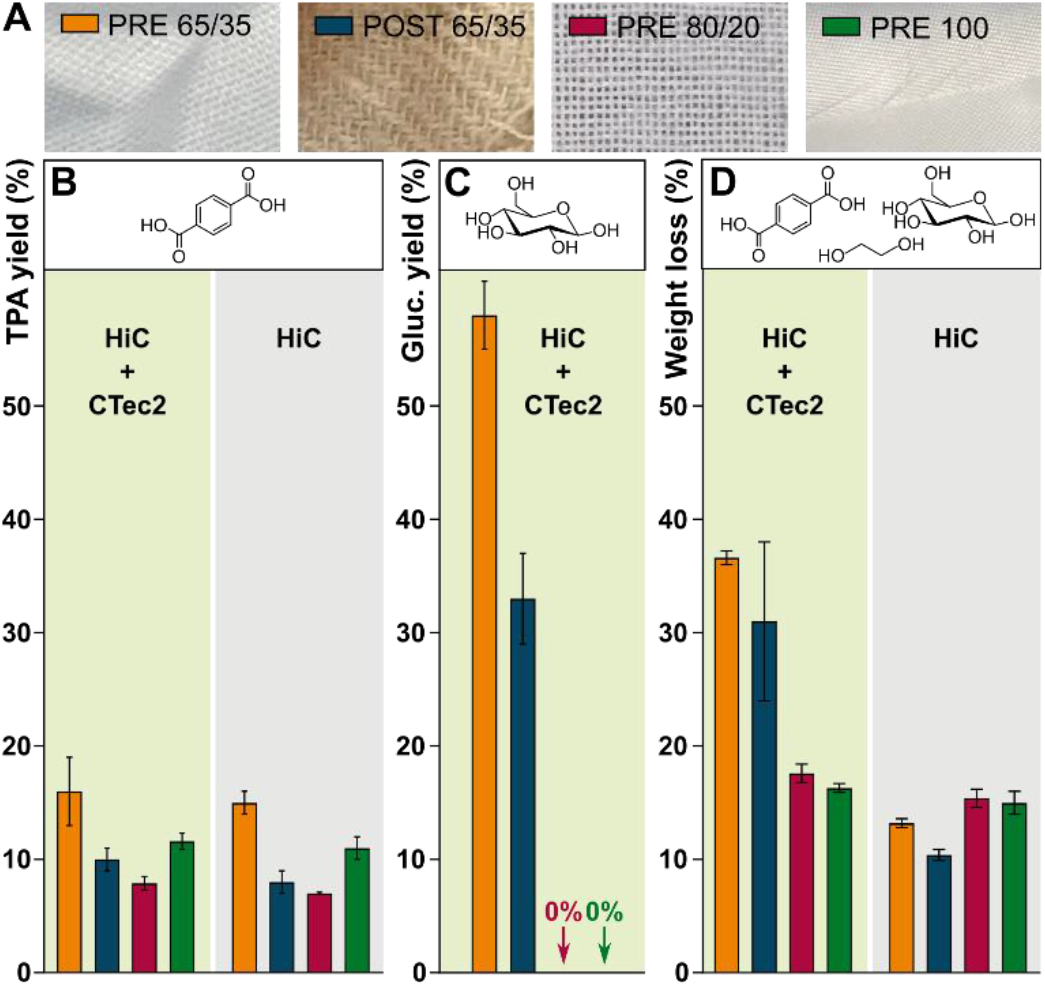
Influence of the textile type (see Table 1 for details; appearance of the respective intact textiles shown in A) and enzyme(s) used on the yield of TPA (B), yield of glucose (C), and total weight loss (D), following RAging reactions (data from Tables S3–S6). Textiles were powdered by pre-milling either for 40 minutes (PRE 65/35, PRE 80/20, PRE 100) or 30 minutes (POST 65/35), prior to enzymatic transformation. Reactions for which HiC enzyme is used together with CTec2 are highlighted with a green background, and those for which HiC is applied alone are highlighted with a grey background.

### RAging experiments with sequential addition of the enzymes

Concluding that CTec2 is inhibited by components of the commercial HiC solution and knowing from previous work^31^ that PET hydrolysis converges due to gradual inactivation of the HiC enzyme, a multi-round experimental design with sequential enzyme addition was developed to maximize the hydrolysis of the **PRE 65/35** textile. Three RAging cycles with CTec2 only, leading to 83 ± 4% yield of glucose were subsequently followed by rounds of PET hydrolysis, where a fresh aliquot of dialyzed and lyophilized HiC powder (lyophilized in order to keep the *η* value unchanged) was added after every 5 RAging cycles, leading to an overall TPA yield of 28 ± 2% after 5 rounds of 5 cycles (total of 25 days, Table S8, entry 1). Importantly, HPLC analysis of the crude products of PET-hydrolysis at the end of the 5-round reaction shows that TPA is produced with 50-fold selectivity over MHET and isolated in 98% purity (Figure S11).

Since the dialyzed and lyophilized HiC powder used above had slightly lower activity than the commercial solution (Table S3, entry 8 *vs* entry 4), an alternative multi-round experiment was designed where fresh HiC aliquots from the commercial solution were added while keeping the liquid-to-solid ratio constant at *η* = 1.5 μL mg^−1^. For this, the first 3 cycles with cellulases were followed by isolation of the glucose product (68.6 ± 0.2% yield of glucose; weight loss of 31 ± 1%), drying the remaining material and restarting the RAging process with addition of the commercial HiC solution. After every 5 cycles, the TPA was extracted by washing, and the remaining solids dried, before adding a fresh aliquot of HiC and restarting the reaction. This allowed us to reach an overall TPA yield of 30 ± 2% and a total of 62.4 ± 0.4% weight loss in 5 rounds (Table S8, entry 2). Furthermore, such a sequential enzymatic approach has the additional advantage of facilitating separation of the glucose and TPA products.

## Conclusions

High-quality recycled PET is highly priced since it is currently produced mostly from clear bottles (a relatively small feedstock) and not available in high enough quantity to meet the rising demand from companies aiming to include recycled material in their products. In fact, the recycled PET feedstock has been shown to be the dominant factor in determining the profitability of new recycling technologies, e.g. enzymatic recycling of PET.^29^ Therefore it is important to consider sources of PET waste thus far not recycled, such as textiles, mixed fibers and layered packaging, when developing new recycling capabilities to meet the need for recycled PET. Interestingly, comparison of TPA production from petrol to that from enzymatic PET depolymerization revealed a 69% lower energy requirement and 17% lower greenhouse gas emissions for the latter, thus encouraging the development of biocatalytic recycling processes.^29^ Besides factors like the feedstock cost and plant size, solid loading and depolymerization extent (not the enzyme cost or loading) were identified as the dominant process factors affecting the profitability of enzymatic PET recycling.^29^ The work presented herein builds towards developing low-waste and environmentally benign^10^ solid-state enzymatic methods,^31–33,42–50^ *i*.*e*. at highest achievable solid loading, for the depolymerization of PET and PET/cotton textiles. While the yield of the TPA monomers from different pre- and post-consumer textiles remains modest (up to 30 ± 2% with a multi-round approach with fresh enzyme additions), compared to glucose yield achieved from cotton (up to 83 ± 4%), this work uniquely addresses the direct enzymatic hydrolysis of highly crystalline PET textiles. Validated here by commercial HiC enzyme, these results could be further enhanced with the use of engineered enzymes. Furthermore, together with previously published results,^31^ this work shows that mechanoenzymatic PET hydrolysis by HiC is unaffected by common contaminants in PET recycling streams, such as polypropylene, colorants, and cotton, and is also not hindered by simultaneous saccharification of cotton in the solid state.

## Author Contributions

**S.K**. Conceptualization, Investigation, Writing – Original Draft, Visualization. **J.A**. Investigation, Validation, Writing – Review & Editing, **T.H.B**. Investigation (Optical microscopy). **J.P.D.T**. Methdology (HPLC method development and HiC enzyme dialysis). **K.A**. Writing – Review & Editing, Project administration, Funding Acquisition. **T.F**. Writing – Review & Editing, Project administration, Funding Acquisition.

## Acknowledgements

We thank the Natural Sciences and Engineering Research Council of Canada (NSERC) for grants RGPIN-2017-06467 and SMFSU 507347-17, the Fond de Recherche du Québec Nature et Technologie (FRQNT) for grant PR 254169, and the Centre in Green Chemistry and Catalysis for funding (FRQNT-2020-RS4-265155-CCVC). S.K. acknowledges funding from the European Union’s Horizon 2020 research and innovation program under the Marie Skłodowska-Curie grant agreement No 101027061.

