## Supporting Information for "Solid-state enzymatic hydrolysis of mixed PET-cotton textiles"

### Materials

The textile samples of mixed PET/cotton (65%/35% and 80%/20%) and PET (100%) were purchased from Fabricville, Canada. The post-consumer PET/cotton (65%/35%) and PET (100%) textile samples were, respectively, a beige scarf provided by Dr. Kaabel and dark blue children's clothing provided by Dr. Auclair. All textile samples were washed with a dilute detergent solution (sodium dodecyl sulfate), rinsed thoroughly with MilliQ water and dried overnight at 55°C in a ventilated oven. MilliQ water was from a Millipore MilliQ system, with a specific resistivity of 18.2 MΩ·cm at 25°C. All solvents used for HPLC analysis (MeCN, MeOH, DMSO, formic acid) were purchased from Millipore Sigma (Oakville, ON, Canada) and Thermo Fisher Scientific (Waltham, MA, US) in HPLC grade. The sodium phosphate buffer (NaPi) was prepared from $NaH_2PO_4$ and $Na_2HPO_4$ (Chem-Impex; Wood Dale, IL, US), using MilliQ water and adjusting the pH of the buffer to 7.3 with HCl (Thermo Fisher Scientific; Waltham, MA, US). The Novozym® 51032 cutinase was expressed in an *Aspergillus* microorganism, for which the source organism is reported as *Humicola insolens* (HiC), was purchased from Strem Chemicals, Inc. (Newburyport, MA, US). The cellulases blend CTec2® (SAE0020) was purchased from Sigma-Aldrich. The enzymes were stored at 4°C. The protein content and the enzyme activity were determined as described in section ***Protein/enzyme characterization***. Multi-round reactions, which included the replenishment of HiC, were carried out with dialyzed and lyophilized enzyme (see section ***Dialysis and lyophilization of the commercial Novozym® 51032 cutinase solution***). The protein content of the resulting enzyme powder was determined using the Bradford assay, and was 9.5% w/w. The Glucose (HK) Assay Reagent (G3293) was purchased from Millipore Sigma and reconstituted in MilliQ water before use. The reconstituted reagent was stored at 4°C for up to 4 weeks. Standards for terephthalic acid (TPA), bis(2-hydroxyethyl) terephthalate (BHET) and glucose were purchased from Millipore Sigma (Oakville, ON, Canada). Mono-2-hydroxyethyl terephthalate (MHET) was synthesized from BHET according to a previously described method.[1]

### Methods

#### *Statistical analysis*

Unless otherwise mentioned, all reactions were performed in triplicate and the error bars in graphs represent the standard deviations.

#### *Equipment*

Mettler Toledo AB135-S/FACT DualRange analytical balance (linearity 0.2 mg, readability 0.01 mg/0.1 mg), Mettler Toledo XP105 DeltaRange analytical balance (linearity 0.15 mg, readability 0.01 mg/0.1 mg), and Gilson and VWR calibrated pipettors were used for measuring out the reactants. Ball milling was carried out using a FormTech Scientific FTS1000 shaker mill, set at a frequency of 30 Hz. Stainless steel

SmartSnap™ jars (15 and 30 mL) or 15 mL unsleeved PTFE jars, all provided by FormTech Scientific, were used as the reaction vessels, charged with either stainless steel or $ZrO_2$ balls, respectively. Aging steps at 55°C were carried out in a Fisher Scientific Isotemp Oven equipped with humidity box saturated with water vapors. Branson 2510 ultrasound bath and Thermo Scientific Legend Micro 21 Centrifuge were used in preparing the samples for HPLC (TPA, MHET and BHET) and plate reader (glucose) analysis. Hydrolysis products of PET were quantified using a HPLC (1260 Infinity II, Agilent Technologies, Santa Clara, CA, USA) equipped with a quaternary pump, autosampler, multiwavelength UV-Vis detector and a Phenomenex Luna® 5 µm C18(2) 100 Å, 4.6 × 250 mm column. The glucose assay was performed using clear bottom 96-well microtiter plates and recording absorbance at 340 nm on a Molecular Devices SpectraMax i3x microplate reader. Lyophilization was performed with a Labconco FreeZone 1 Liter Benchtop Freeze Dry System. A dynamic scanning calorimeter Discovery 2500 from TA Instruments was used to determine the crystallinity of the pure PET samples (100% polyester textile), and to verify the polyester type (PET) in all textile materials based on the melting temperature. Attenuated total reflectance Fourier transform infrared (ATR-FTIR) spectra were obtained in the 400 $cm^{-1}$ to 4000 $cm^{-1}$ range on a Bruker VERTEX 70 FTIR spectrometer with an integrated Platinum ATR unit. A WITec 300 R confocal Raman microscope, equipped with a $\lambda$ = 532 nm laser probe, DV 401 BV CCD camera, and a WITec UHTS 300 spectrometer, was used to capture the optical microscopy images of the textile materials. Raman spectra were recorded, in an attempt to map the distribution of PET and cotton in the intact and pre-milled **PRE 65/35** textile, however due to the high ratio of PET *vs* cotton in this textile, only the bands corresponding to PET were observable. In fact, Was-Gubala and Machnowski have shown that when using an Ar ion laser ($\lambda$ = 514 nm) to characterize dyes in cotton and viscose textiles, the spectra were dominated by the dye bands (dyes were present in the textile in 0.2 – 3 wt%), while no bands originating from cellulose could be observed. [2]

##### *Pre-treatment of the textiles*

All textile samples were washed with a dilute detergent solution (sodium dodecyl sulfate), rinsed thoroughly with MilliQ water and dried overnight at 55°C in a ventilated oven. When pre-milling was mentioned, the textile samples were powdered by milling at 30 Hz for 20 – 60 minutes, which increases the surface area of the material and avoids inconsistency resulting from non-homogeneous solid reaction mixtures. Thus, 0.6 or 1 g of textile were weighed into a 30 mL stainless steel milling jar, before addition of a 11.7 g stainless steel ball, and milling for 20-60 minutes at 30 Hz.

##### *Crystallinity determination of PET textiles, polymer verification by DSC and polyester content*

Differential scanning calorimetry (DSC) measurements were performed using 2–8 mg of the pre-milled textile samples placed into hermetically sealed aluminum pans, which were then analyzed by heat-cool-heat cycles in the temperature range 0 to 300°C. The degree of crystallinity ($X$c) of the semi-crystalline PET polymers in the pure PET textiles were estimated using the TRIOS software version 4.5.1.42498, based on the first heating scan from 0°C to 300°C at a rate of 10°C $min^{-1}$, according to the equation $X\mathrm{c} = \frac{\Delta H_\mathrm{f} - \Delta H_c}{\Delta H_\mathrm{f}^\mathrm{o}}$, where $\Delta H_\mathrm{f}$ is the enthalpy of fusion, $\Delta H_c$ is the enthalpy of cold crystallization (not exhibited by any of the textile materials) and $\Delta H_\mathrm{f}^\mathrm{o}$ is the enthalpy of fusion of a 100% crystalline PET polymer (140 J $g^{-1}$) at the melting temperature. The enthalpy of fusion $\Delta H_\mathrm{f}$ was determined by integration of the endothermic melting peak, wherein a straight baseline was drawn between the visually determined starting point of melting to the end point.

The polymer type of the textiles labelled as “polyester” was determined from the melting temperature of the second heating cycle, where the previous thermal history of the PET polymers is “erased” and equalized by the first melting and cooling cycle. All studied polyesters melted in the range of 249–256°C corresponding to the melting temperature of PET.

The polyester content of the **PRE 65/35**, **PRE 80/20** and **PRE 100** was measured according to a method adopted from the SFS-EN ISO 1833-1:2020 standard, based on selective hydrolysis of cellulose from binary fiber mixtures and weighing the remaining residue (which is taken as equal to polyester content). Intact washed and dried textile pieces of **PRE 65/35** (195.5 mg), **PRE 80/20** (180.2 mg) and **PRE 100** (142.1 mg) were weighed into round-bottom flasks, and 30 mL of 70% $H_2SO_4$ was added to each. Each reaction was stirred at room temperature for 1 hour, and then quenched by pouring into cold water (on an ice bath). The remaining residue was washed on a filter with water, freeze-dried and weighed. The residue weight was 71% for **PRE 65/35**, 85% for **PRE 80/20**, and 100% for **PRE 100**. While the residue weight is proportional to the label polyester content in each textile, the slightly higher weight of residue compared to the polyester content reported by the producer can arise from non-complete hydrolysis of cellulose or non-fibrous matter (dyes or resins added for water-repellency or crease-resistance) in the textile. Extended acid hydrolysis (2 hours) gave polyester contents of 67% for **PRE 65/35**, 89% for **PRE 80/20**, and 99% for **PRE 100**.

##### *Characterization of the enzymes*

Unless noted otherwise, both enzymes were used as the supplied aqueous solutions, which contain propylene glycol as antifreeze. The protein concentrations of the enzyme solutions were determined by using the Bradford assay and were 6.5 mg $mL^{-1}$ of protein in the Novozym® 51032 cutinase solution and 6.1 mg/ml of protein in the cellulase blend CTec2®.

The activity of the cutinase enzyme was determined by assaying for hydrolysis of bis(2-hydroxyethyl) terephthalate (BHET). This was carried out using solutions containing 0.65 mg $mL^{-1}$ of the enzyme, diluted in 0.1 M sodium phosphate buffer at pH 7.3. This enzyme solution (99 µL) was incubated at 55°C for 10 minutes, after which the reaction was initiated with the addition of 1 µL of a 100 mM BHET stock solution in DMSO to achieve a final concentration of 1 mM BHET, and allowed to shake (250 rpm) at 55°C. The reaction was quenched by adding MeOH (400 µL), followed by sonication and centrifugation, after which the clear supernatant was analyzed by HPLC. The commercial cutinase preparation converted ~99% of the BHET into MHET after only 10 min, with no loss in enzymatic activity over several months.

The activity of the cellulase enzymes was determined by assaying the hydrolysis of Type 20 microcrystalline cellulose (MCC) into glucose at 37°C. The assay was carried out using a 5% (w/v) suspension of MCC in 50 mM sodium acetate buffer (pH 5.0 at 37°C). The commercial CTec2 solution (6.1 mg $mL^{-1}$ protein content) was diluted to 0.04 mg $mL^{-1}$ in MilliQ water before addition to the reaction mixture. The assay was started by incubating 400 µL of the MCC suspension at 37°C. Then 100 µL of the 0.04 mg $mL^{-1}$ enzyme solution (total enzyme dilution factor from the commercial solution df = 750) was added and the mixture was allowed to shake (250 rpm) at 37°C for 120 min. The reaction was stopped by placing the assay sample on ice, after which it was centrifuged at 4000 rpm and the clear supernatant was assayed for glucose content. The activity of CTec2 was measured as 2406 U $mL^{-1}$. Variability in the activity of the purchased CTec2 enzymes was noticed, with the second batch showing 1828 U $mL^{-1}$ activity (76% of the original batch). Experiments where the second batch of CTec2 were used (post-consumer textile hydrolysis experiments, Table S2 entry 3 and Table S6 entry 2) were carried out with the same amount of enzyme preparation as with the original batch, and therefore the lower glucose yield of these two reactions were affected by the lower CTec2 activity.

##### *Method for milling + aging reactions*

In a typical reaction, pre-milled textile powder (200 mg) was weighed into a 15 mL stainless steel jar with a 3.5–4 g stainless steel ball, to which the commercial enzyme preparations (either 200 µL HiC or 200 µL

HiC + 20 µL CTec2) and buffer (100 µL) were added, bringing the total liquid-to-solid ratio to 1.5 µL $mg^{-1}$, corresponding to a solid loading of 40% w/w for the reaction mixture. The CTec2 commercial enzyme preparation contains a high concentration of inherent glucose (*ca*. 20% w/w, see section ***Enzyme inherent glucose content determination***) and the volume of this enzyme solution was therefore not counted as liquid in the calculation of the total liquid-to-solid ratio. Reactions starting from intact textile (no pre-milling) were performed with 0.5 $cm^2$ hand-cut textile pieces, instead of the pre-milled material. The added 200 µL of Novozym® 51032 cutinase solution (6.5 mg $mL^{-1}$ of protein) corresponds to 0.6% w/w of HiC enzyme respective to the textile mass, while the 20 µL of CTec2® cellulases blend (70 mg $mL^{-1}$) corresponds to 0.7% w/w of enzyme relative to the textile mass, assuming that the commercial preparations contained no other proteins. The milling jars were then closed and set to mill at 30 Hz for 5 or 30 minutes. The resulting paste-like uniform solids were aliquoted (20–50 mg) into 1.5 mL Eppendorf tubes for aging at 55°C, affording analysis samples for HPLC and glucose assay at different time points. Specific reaction conditions, together with the hydrolysis yields are compiled into **Table S1** (PRE 65/35 textile) and **Table S2** (post-consumer textiles).

##### *Quantification of PET hydrolysis products by HPLC*

TPA, MHET, and BHET standards for the calibration curves were prepared from 10 mg $mL^{-1}$ stock solutions in DMSO, by diluting to 200, 100, 50, and 25 µg $mL^{-1}$ in MeOH. All calibration standards were prepared in triplicate. Separation of TPA, MHET and BHET was achieved on a Phenomenex Luna® 5 µm C18(2) 100 Å, 4.6 × 250 mm column. An older column, with poorer separation was used with a 40-minute gradient elution from 99(A):1(B) to 60(A):40(B), where A is 0.1% formic acid in water and B is acetonitrile, the injection volume was set to 10 µL, the flow rate used was 0.6 mL $min^{-1}$ and UV detection was set at 240 nm. The retention times for TPA, MHET and BHET were 26.9, 30.1 and 31.7 min, respectively. A new identical column was purchased, which allowed to shorten the elution method to a 21-minute gradient elution from 99(A):1(B) to 65(A):35(B), with a flow rate of 1 mL $min^{-1}$. The retention times for TPA, MHET and BHET with this column were 16.2, 18.1 and 19.1 min, respectively. The calibration curves of TPA, MHET and BHET were constructed by plotting the areas under the corresponding peaks *vs* concentration of the standard solutions (25 – 200 µg $mL^{-1}$). The calibration curves were used to interpolate the concentration of hydrolysis products in reaction mixtures.

The PET hydrolysis products were extracted from the reaction mixture by adding a corresponding amount of DMSO to the reaction aliquot, to achieve a 20 mg $mL^{-1}$ suspension, which was then sonicated for 1–5 minutes at room temperature to ensure the dissolution of all hydrolysis products. It is important to note, that no hydrolysis of PET or BHET into MHET or TPA takes place during sonication in DMSO (assayed up to 30 minutes), therefore the composition of the analyzed products (TPA, MHET, BHET) is unaffected by sample preparation. The suspension was then centrifuged at 21 000 × *g* for 5 minutes to separate the remaining insoluble solids (containing PET) from the solution and the supernatant was diluted 10- to 40-fold in MeOH to prepare HPLC samples of concentrations within the calibrated range. All HPLC samples were syringe filtered using 0.22 µm nylon filters (Chromspec, Brockville, ON, Canada) prior to analysis.

The yield of TPA (%) is calculated by dividing the experimentally determined yield of TPA (mg), extrapolated from the content of TPA in solids (determined by HPLC), with the theoretical yield at 100% conversion. Since PET-cotton mixed textiles were analyzed in this study, the theoretical yield considers the PET content and is calculated by approximating the PET polymer as an infinite linear chain of condensed TPA and ethylene glycol (EG). For example, complete hydrolysis of 200 mg (1.04 mmol) of PET would yield 173 mg (1.04 mmol) of TPA and 64.6 mg (1.04 mmol) of ethylene glycol, assuming a 100% pure PET material. The TPA yield achieved from mixed material textiles considers the PET content. For example, complete hydrolysis of 200 mg of 65% PET textile (0.68 mmol) would yield 112.7 mg (0.68 mmol) of TPA and 42 mg (0.68 mmol) of ethylene glycol.

##### *Determination of glucose yield by the glucose assay*

A commercial glucose assay reagent was used to quantify glucose in the reaction mixtures. The aliquot of the reaction mixture to be analyzed was suspended at 10 mg $mL^{-1}$ in cold water, closed tightly and quickly placed in boiling water to denature the enzymes. After 30 minutes of boiling, the sample was sonicated for 5 minutes to ensure the dissolution of glucose from solid aggregates, and the suspension was centrifuged at 21 000 × *g* for 5 minutes. An aliquot of the clear supernatant (2 µL) was mixed with the glucose assay reagent (150 µL) in a clear-bottom 96-well plate and incubated at room temperature. The absorption was monitored at 340 nm until the reading plateaued. Glucose calibration standards (0 – 10 mg $mL^{-1}$) were measured on each occasion alongside sample measurements, due to the gradual inactivation of the Glucose Assay Reagent. The resulting calibration curve, which relates the glucose concentration to absorbance at 340 nm was used to interpolate the glucose concentration in the reaction samples.

The yield of glucose (%) is calculated by dividing the experimentally determined yield of glucose (mg) with the theoretical yield at 100% conversion. Since PET-cotton mixed textiles were analyzed in this study, the theoretical yield considers the cotton content and is calculated by approximating the cellulose in cotton to infinite linear chains of condensed glucose. The experimentally determined yield of glucose is extrapolated from the content of glucose in solids as determined by the glucose assay, with enzyme-inherent glucose subtracted (see further details in section ***Enzyme inherent glucose content determination***).

##### *Enzyme inherent glucose content determination*

Previous experience from our group has shown that commercial enzyme preparations can inherently contain some glucose, and/or carbohydrates from which glucose is produced during aging.[3] This glucose was quantified in this work at two timepoints: straight after milling and after 3 days of subsequent aging for non-cellulose containing model reaction mixtures. Model reaction mixtures were created using a commercial pure PET sample (Goodfellow, contains no glucose), ensuring that all glucose detected in the samples originates from the enzyme mixtures. The reaction mixtures were prepared as described in ***Method for milling + aging reactions***, taking assay samples at 10, 20, 30 and 50 mg/ml, to ensure that the glucose content is detectable at the used assay conditions. Samples taken were treated and analyzed as described in section ***Determination of glucose yield by the glucose assay***. Based on all samples ($n$ = 12 for each timepoint) the solids were found to contain 2.6 ± 0.2% glucose after milling and 2.9 ± 0.1% glucose after subsequent aging for 3 days, which shows that the commercial enzyme preparation indeed contains glucose (*ca*. 20% w/w) and that the amount of glucose slightly increases during aging. Glucose assay of the hydrolysis outcome of 100% PET materials with HiC enzyme alone showed that no glucose is present in these solids, therefore all enzyme-inherent glucose is introduced with the CTec2 solution. The amount of inherent glucose in milled and aged reaction mixtures was accounted for (subtracted) in the calculation of the hydrolysis yield of cellulose.

##### *Method for the Reactive Aging reactions (RAging)*

In a typical RAging reaction, pre-milled textile powder (200 mg) was weighed into a 15 mL unsleeved PTFE jar, charged with a single 10 mm $ZrO_2$ ball (3.5 g), to which the commercial enzyme preparation(s) (either 200 µL HiC or 200 µL HiC + 20 µL CTec2) and buffer (100 µL) were added, bringing the total liquid-to-solid ratio to 1.5 µL $mg^{-1}$, corresponding to a solid loading of 40% w/w for the reaction mixture. The CTec2 commercial enzyme preparation contains a high concentration of inherent glucose (*ca*. 20% w/w, see section ***Enzyme inherent glucose content determination***) and the volume of this enzyme solution was therefore not counted as liquid in the calculation of the total liquid-to-solid ratio. Reactions at higher liquid-to-solid ratio, 2 µL mg$^{-1}$ (32 % w/w), 2.5 µL mg$^{-1}$ (28 % w/w) and 3 µL mg$^{-1}$ (25 % w/w), were set up identically, except for the addition of 100, 200 or 300 µL of water, respectively. A RAging cycle consisted of ball milling at 30 Hz for 5 minutes, followed by aging at 55°C for 24 hours. The aging duration was chosen based on the kinetic analysis of the HiC enzyme on solid PET substrate.[1] The RAging cycles were repeated 7 times, with an aliquot (10–20 mg) collected after each cycle for analysis of the reaction products by HPLC. Sealing tape proved to be necessary to ensure that the jars remain closed and water evaporation was minimal during the aging step. Specific reaction conditions and variations tested, together with corresponding hydrolysis yields are compiled into **Table S3** (PRE 65/35 textile), **Table S4** (PRE 80/20 textile), **Table S5** (PRE 100 textile) and **Table S6** (POST 65/35 textile).

##### *Dialysis and lyophilization of the commercial Novozym® 51032 cutinase solution*

This protocol was applied for the multi-round RAging reactions described in section ***Method for the multi-round RAging reactions with fresh enzyme additions***. The commercial HiC (estimated MW of 20–30 kDa)[4] solution was diluted to a concentration of 3.25 mg mL$^{-1}$ (from initial 6.5 mg mL$^{-1}$) with 25 mM Tris buffer (pH 7.3) containing 100 mM NaCl, and then placed in a Repligen Spectra/Por® 3 Standard Regenerated Cellulose (RC) Dialysis tube with a MWCO of 3.5 kDa (Spectrum Chemical Mfg. Corp.; New Brunswick, NJ, US) and dialyzed against 1 L of a 25 mM Tris buffer (pH 7.3) containing 100 mM of NaCl, replaced every hour for 10 hours. The content of the dialysis tube was then placed into a 50 mL conical tube, flash-frozen in an acetone and dry ice bath and lyophilized until dry. The dry powders were stored in tightly closed containers at 4°C. The protein content of the lyophilized powder was determined to be 9.5% w/w by using the Bradford assay.

##### *Method for the multi-round RAging reactions with fresh enzyme additions*

In multi-round RAging reactions, 600 mg of pre-milled PET-cotton textile powder (65% PET, 35% cotton) was weighed into a 15 mL unsleeved PTFE jar, charged with a single 10 mm $ZrO_2$ ball (3.5 g). **1st round:** Since CTec2® proved more efficient when acting alone, 60 µL of CTec2® was pre-mixed with 300 µL of the buffer and 540 µL of water (this substitutes the amount of water generally introduced with the HiC solution) and added to the reaction jar. The jars were closed with sealing tape to minimize water evaporation. Three cycles of RAging were performed, each consisting of 5 minutes of milling at 30 Hz followed by aging at 55°C for 24 hours, after which an aliquot of the reaction mixture (15 mg) was taken for determining the glucose yield. **2nd, 3rd, 4th and 5th round:** The round starts with the addition of 26.5 mg of dialyzed and lyophilized HiC powder (see section ***Dialysis and lyophilization of the commercial Novozym® 51032 cutinase solution***) and 27 µL of water to the jar, followed by five cycles of RAging. Sample aliquots (10 – 30 mg) for determining the PET and cotton hydrolysis yields were taken at the end of each round, and the results are compiled in **Table S7**.

Alternatively, a protocol for a multi-round RAging reaction with end-of-round product wash-out was also developed. Similar to above, 600 mg of pre-milled PET-cotton textile powder (65% PET, 35% cotton) was weighed into a 15 mL unsleeved PTFE jar, charged with a single 10 mm $ZrO_2$ ball (3.5 g) and 60 µL of CTec2® pre-mixed with 300 µL of the buffer and 540 µL of water. **1st round:** Three cycles of RAging were performed, each consisting of 5 minutes of milling at 30 Hz followed by aging at 55°C for 24 hours. After this, an aliquot of the reaction mixture (15 mg) was taken for glucose analysis and the remaining solids were washed in a 50 mL conical polypropylene tube three times as follows: the reaction mixture was suspended in 20 mL of cold water, vortexed thoroughly and centrifuged at 10 000 × *g* for 25 minutes. The clear supernatant was removed and discarded after each wash. The remaining solid was frozen with liquid nitrogen and lyophilized. The weight loss of the solids resulting from the 1st round was 31 ± 1%. **2nd, 3rd, 4th and 5th round:** Each round starts with loading the dry remaining solids into the 15 mL unsleeved PTFE jar, together with a corresponding amount of the commercial solution of HiC (0.65% $w_{protein}/w_{solid}$) and buffer, to maintain a liquid-to-solid ratio of 1.5 µL $mg^{-1}$. The jars were charged with a single 10 mm $ZrO_2$ ball (3.5 g) and sealed with tape, followed by five cycles of RAging. Sample aliquots (10 – 30 mg) for quantifying the PET and cotton hydrolysis products were taken at the end of every round (here 1 round = 5 cycles of RAging), before products were washed out. After this, the PET hydrolysis products were removed by washing the reaction mixture in a 15 mL conical tube with 0.3 M $Na_2CO_3$, vortexing and centrifuging at 10,000 × g for 5 min. The supernatant containing the soluble sodium terephthalate was removed. The remaining solid was washed with MilliQ water (3 × 3 mL), until neutral pH was reached, frozen with liquid nitrogen and lyophilized. The hydrolysis yields after each round are compiled into **Table S7**.

##### *Determining the weight loss of textiles after mechanoenzymatic hydrolytic reactions*

An aliquot of 50-60 mg of the paste-like reaction mixture was taken from the RAging jars at the end of the reaction, dried by lyophilization to determine the precise dry mass of the sample ($m_{dry}$), and then washed 2 × 1 mL of $H_2O$ and 4 × 1 mL MeOH, to remove the hydrolysis products. Between the washes the suspension was sonicated, to ensure the dissolution of all soluble hydrolysis products, and centrifuged at 21 000 × $g$ for 5 minutes to allow separation of the liquid. The sample was again dried by lyophilization, revealing the weight of the remaining textile material ($m_{washed}$). The weight loss (%) was determined from: $Weight\ loss\ (\%) = \frac{m_{dry} - m_{washed}}{m_{dry}} \cdot 100\%$. The dry mass ($m_{dry}$) of the reactions which included the CTec2 enzyme contain 2.9 wt% of the enzyme-inherent glucose (see section ***Enzyme inherent glucose content determination***), which was subtracted from the corresponding $m_{dry}$ prior to calculating the weight loss, so that the latter would represent only the weight loss due to hydrolysis.

Based on the experimentally determined content of hydrolysis products in solids, an expected weight loss was also calculated. This was done by calculating the sum weight of TPA (content in solids determined by HPLC), glucose (content in solids determined by the glucose assay), and ethylene glycol (calculated assuming 1:1 molar ratio to TPA) which are produced in the reaction and subtracting these from the amount of initial textile substrate. The estimated weight loss for **PRE 65/35** and **PRE 100** matched well with the experimental weight loss (Figure 3C), with the experimentally determined weight loss falling within ±10% of the calculated weight loss. However, the experimentally determined weight loss of **PRE 80/20** was *ca.* 2.0 times higher compared to the respective estimated weight loss.

#### Supplementary Figures

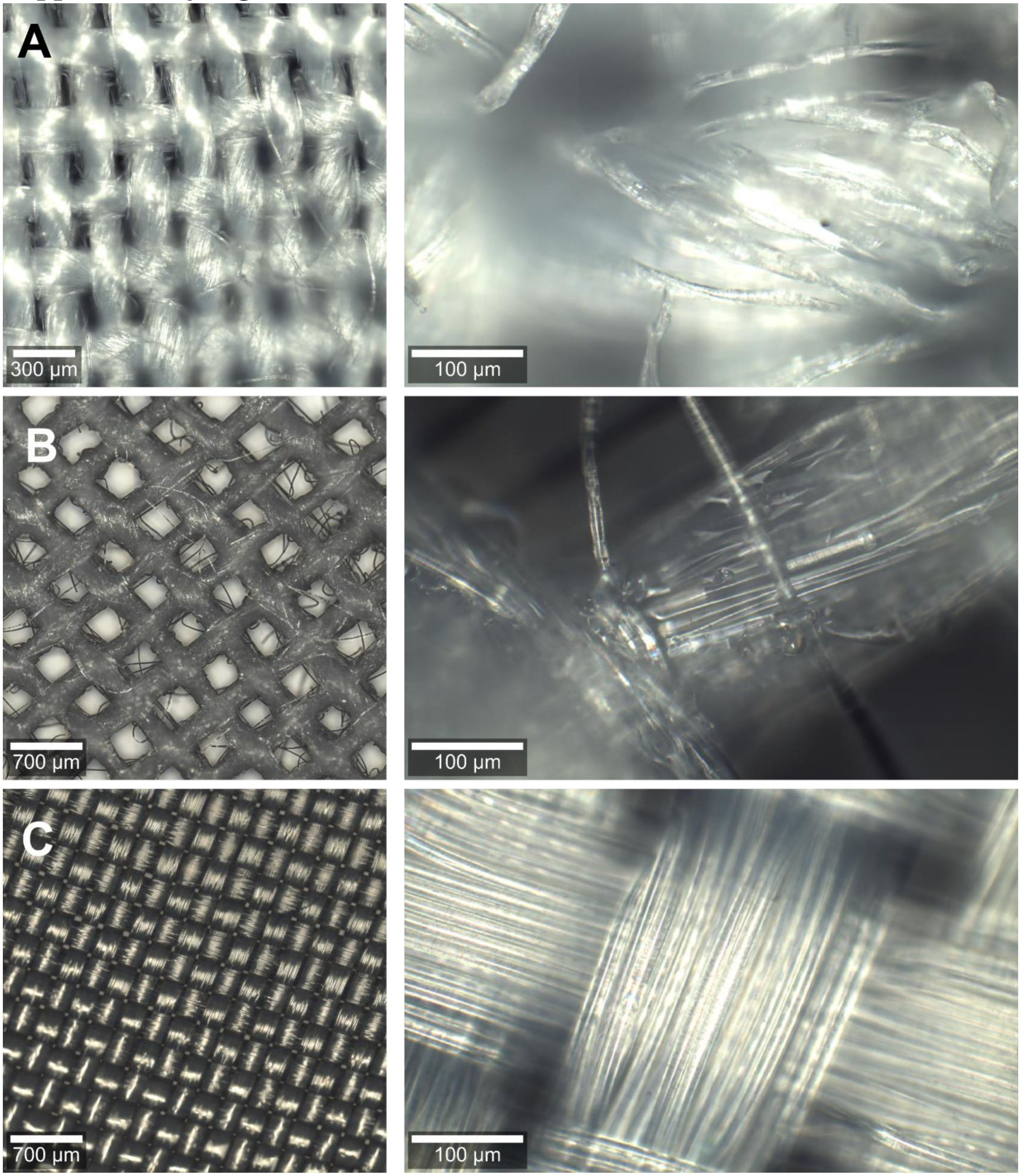

***Figure S1.***
Optical microscopy images of the studied pre-consumer textiles: A) 65% PET, 35% cotton broadcloth textile (**PRE 65/35**); B) 80% PET, 20% cotton buckram textile (**PRE 80/20**); C) 100% PET lining textile (**PRE 100**).

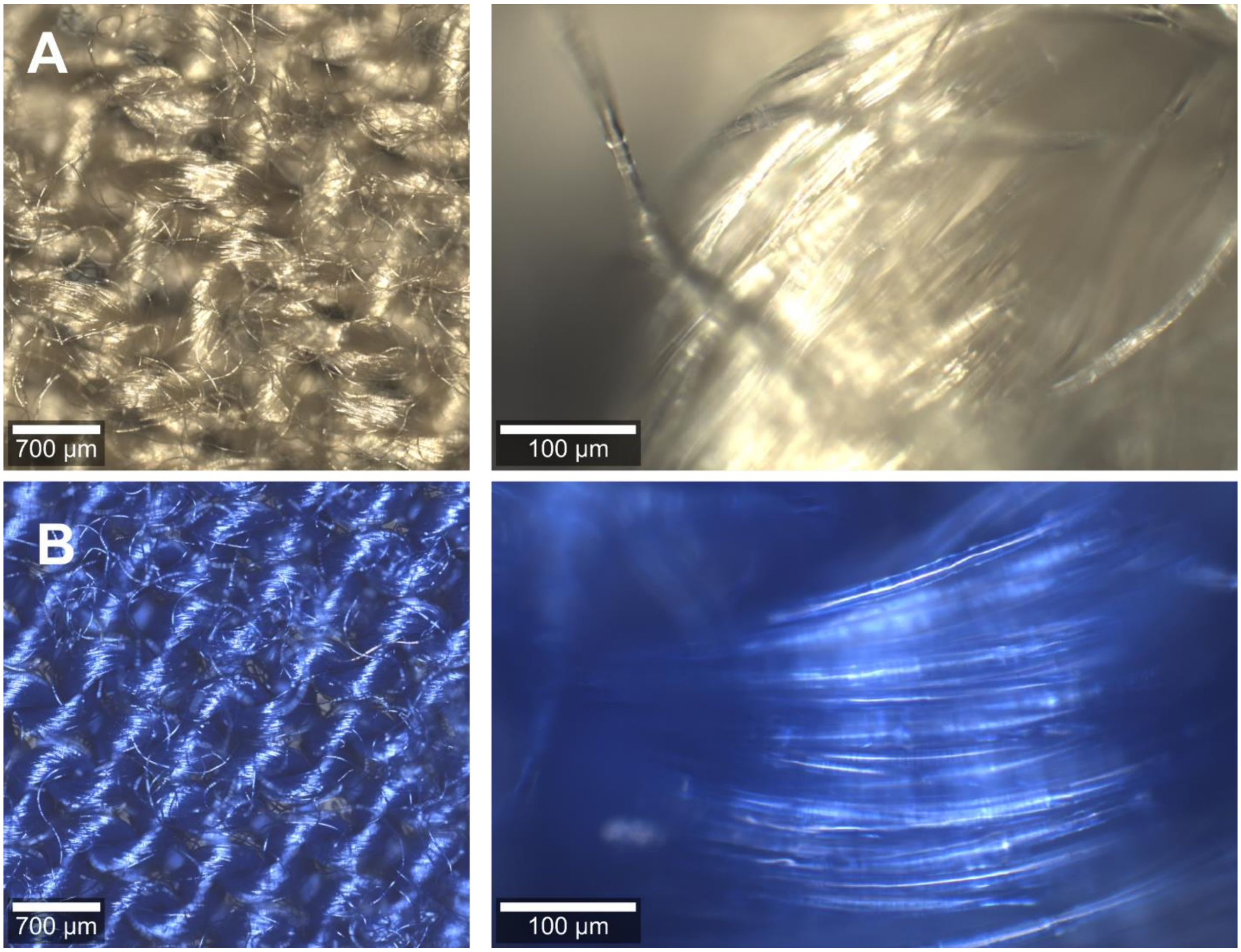

***Figure S2.***
Optical microscopy images of the studied post-consumer textiles: A) beige 65% PET, 35% cotton textile (**POST 65/35**); B) dark blue 100% PET textile (**POST 100**).

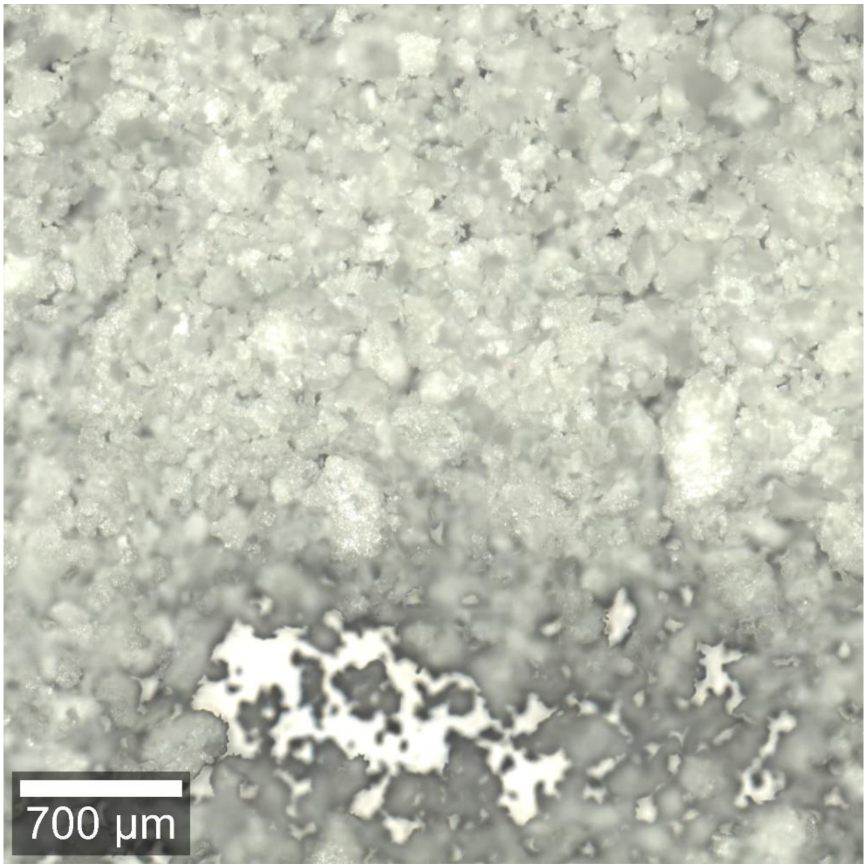

***Figure S3.***
Optical microscopy image of the pre-milled (40 minutes, 30 Hz) 65% PET 35% cotton textile (**PRE 65/35**), which shows that the fibers are completely disrupted by milling.

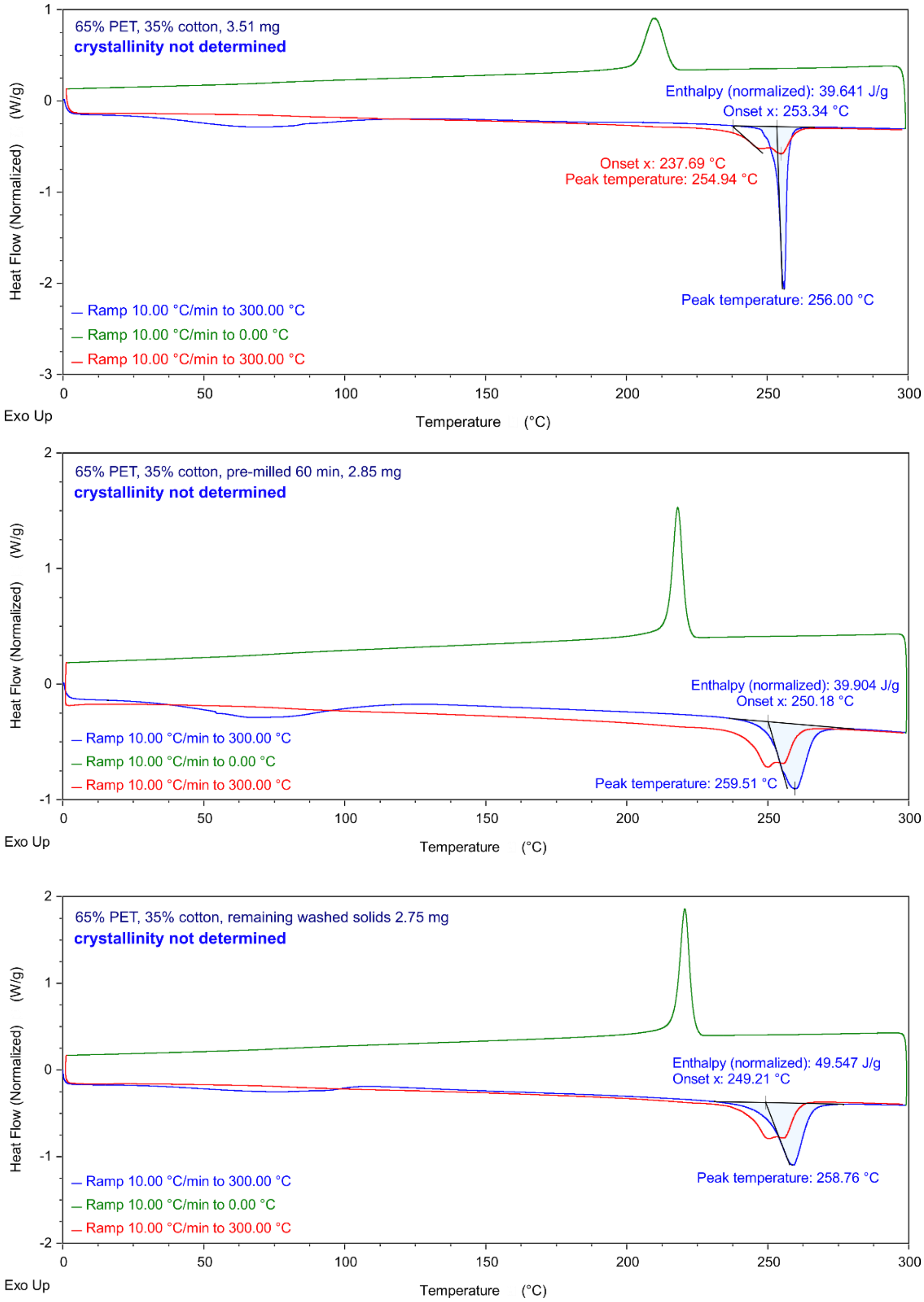

***Figure S4.***

Differential scanning calorimetry heat (blue) – cool (green) – heat (red) scans in the range 0°C to 300°C at a rate 10°C $min^{-1}$ on the **PRE 65/35** pre-consumer white 65% PET and 35% cotton textile as purchased (**top scan**), pre-milled 60 minutes (**middle scan**) and post-reaction remaining washed solids (**bottom scan**). The melting temperature of the material 255°C (top scan, second heating cycle, shown in red) verifies the polymer type as PET. While % crystallinity could not be determined from the melting enthalpy of PET, since this is a mixed PET/cotton material, the relative comparison of as purchased (top scan) and pre-milled material (middle scan) show that pre-milling does not change the crystallinity of PET (melting enthalpy is proportional to crystallinity if PET/cotton ratio is kept constant). Significantly increased melting enthalpy for the post-reaction remaining solids (bottom scan, washed material from Table S3, entry 7) indicates that the ratio of PET is increased after the hydrolysis reaction (see Figure S8).

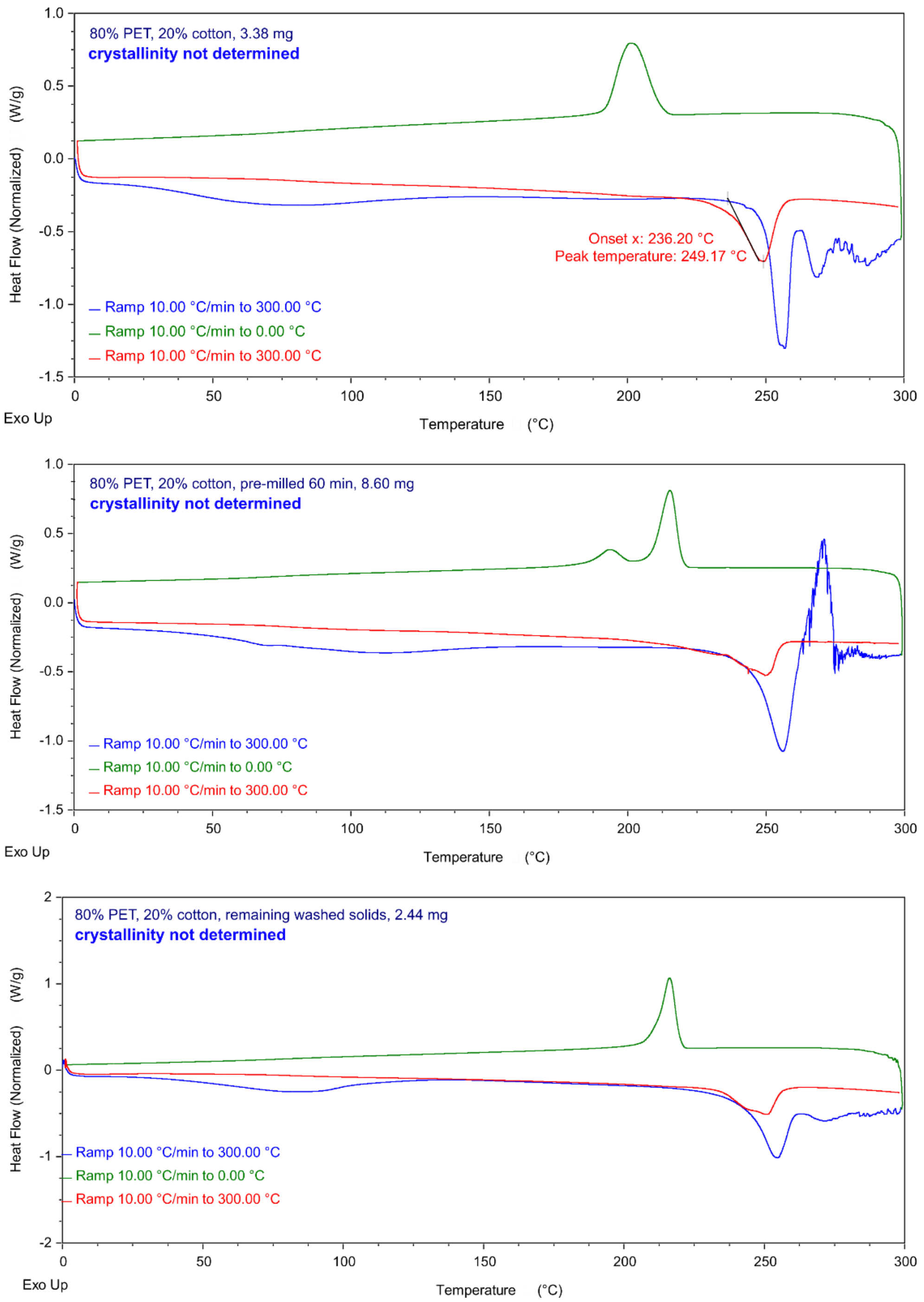

***Figure S5.***

Differential scanning calorimetry heat (blue) – cool (green) – heat (red) scans in the range 0°C to 300°C at a rate 10°C min$^{-1}$ on the **PRE 80/20** pre-consumer white 80% PET and 20% cotton textile as purchased (**top scan**), pre-milled 60 minutes (**middle scan**) and post-reaction remaining washed solids (**bottom scan**). The melting temperature of the material 249°C (top scan, second heating cycle, shown in red) verifies the polymer type as PET (mp range 240–265°C). % crystallinity could not be determined from the melting enthalpy of PET, since this is a mixed PET/cotton material. The first heating scans of this material show additional endothermic events after the PET melting, which may be due to the particular thermal history of PET in this fabric. PET crystallization and thus melting behavior is known to depend on the cooling rate used in the processing of the molten polymer,[5] and PET yarn crystallinity is additionally influenced by the texturing (high temperature) conditions.[6] Therefore, multiple melting peaks reflect the processing of the PET fibers, and this is why only the second heating scan was used for polymer identification.

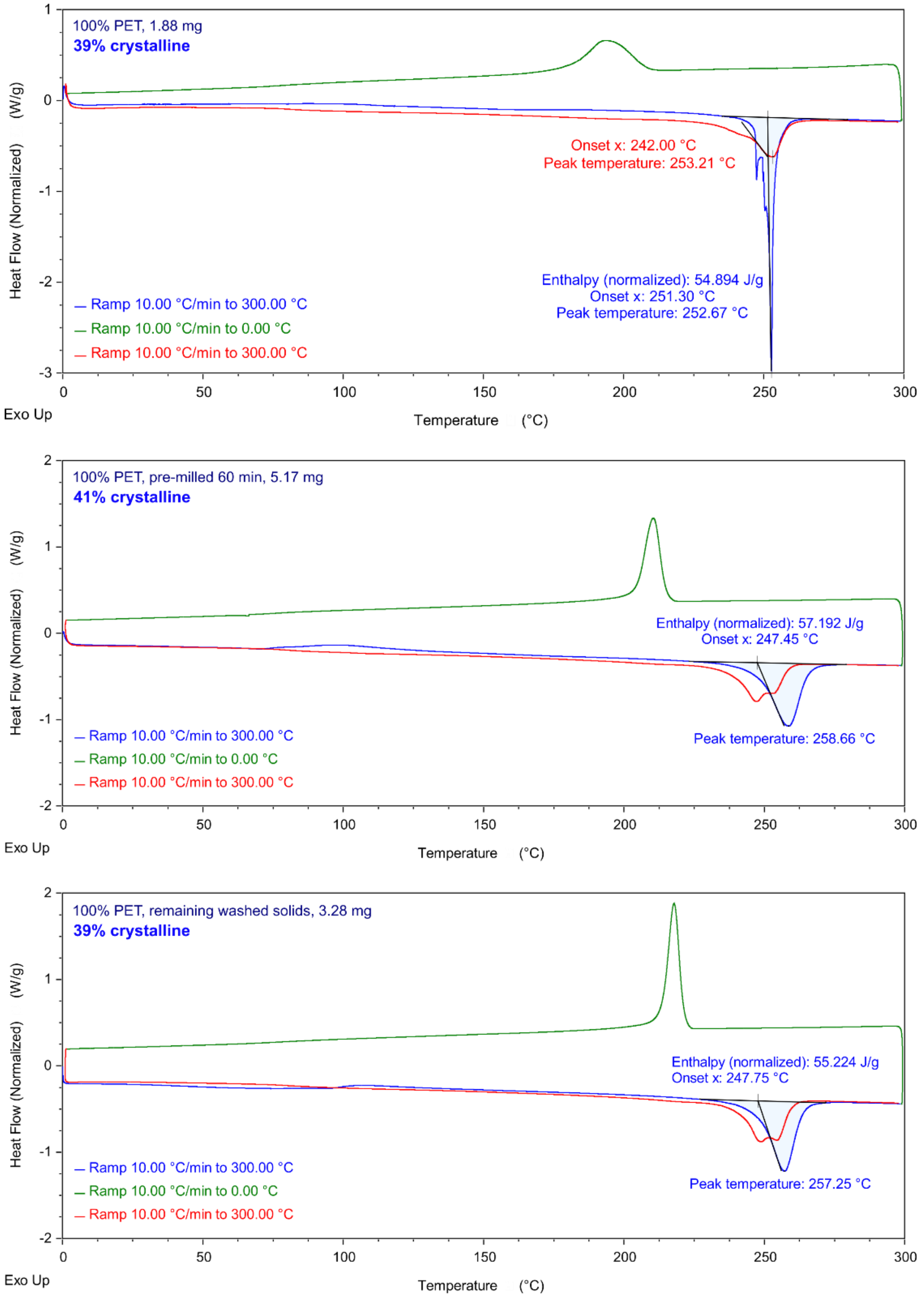

***Figure S6.***

Differential scanning calorimetry heat (blue) – cool (green) – heat (red) scans in the range 0°C to 300°C at a rate 10°C min$^{-1}$ on the **PRE 100** pre-consumer white 100% PET textile as purchased (**top scan**), pre-milled 60 minutes (**middle scan**) and post-reaction remaining washed solids (**bottom scan**). The melting temperature of the material 253°C (top scan, second heating cycle, shown in red) verifies the polymer type as PET (mp range 240–265°C).

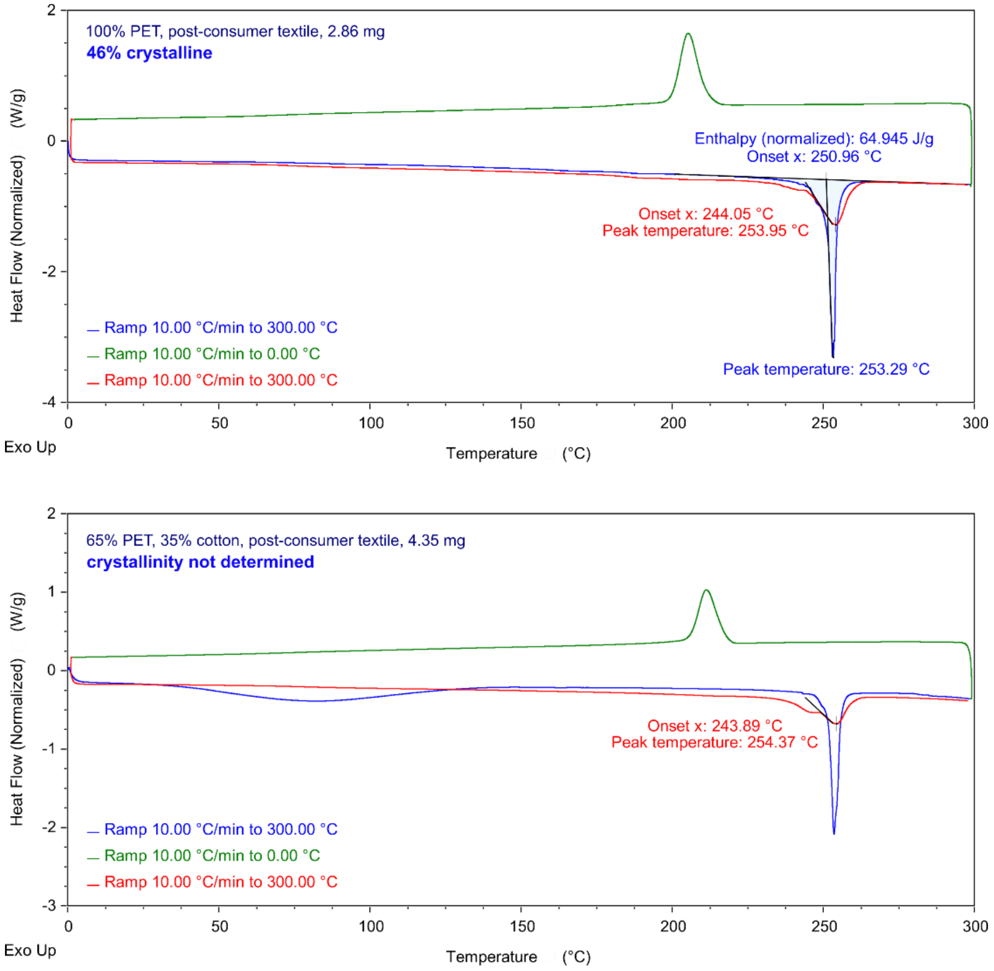

***Figure S7.***

Differential scanning calorimetry heat (blue) – cool (green) – heat (red) scans in the range 0°C to 300°C at a rate 10°C min$^{-1}$ on the post-consumer textiles: 100% PET dark blue textile (**POST 100**, **top scan**) and the 65% PET and 35% cotton beige fabric (**POST 65/35**, **bottom scan**). The melting temperature of the material 254°C (second heating cycle on both scans, shown in red) verifies the polymer type as PET (mp range 240–265°C).

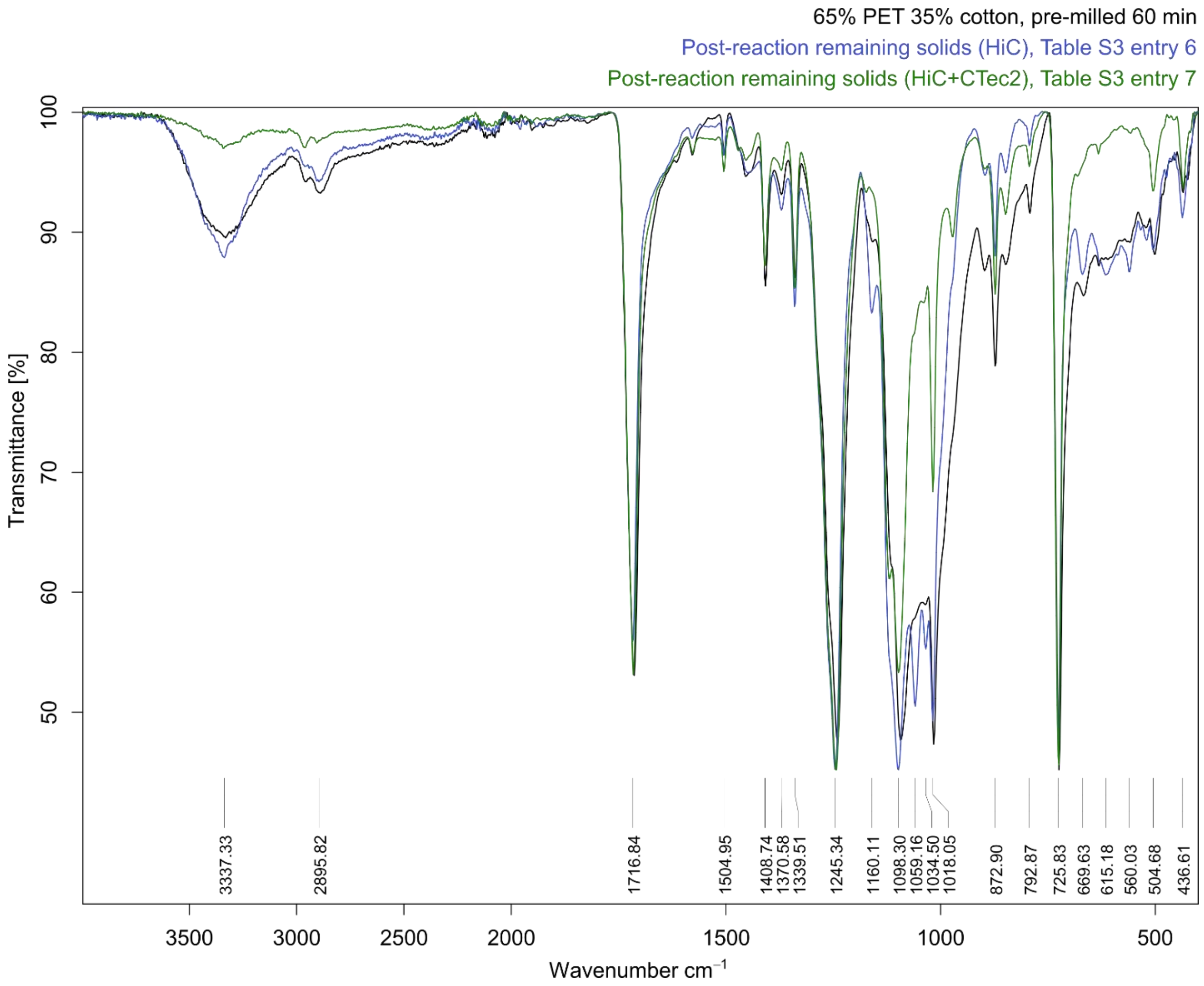

***Figure S8.***

FTIR spectra of the **PRE 65/35** pre-consumer white 65% PET and 35% cotton textile before (pre-milled textile, spectra shown in black) and after the mechanoenzymatic reaction (blue and green), shown in full recorded range. The post-hydrolysis washed materials show the difference of applying HiC enzyme alone (blue) or together with CTec2 cellulases (green), which lead to 11.0 ± 0.9% and 39 ± 2% weight loss, respectively. Extensive hydrolysis of cotton by CTec2 cellulases is evident in the spectra shown in green, from the significant reduction of the absorbance bands characteristic to cotton at 3337, 2896 and 1160 $cm^{-1}$ (polysaccharide O-H and C-H stretching vibrations and C-O-C asymmetrical stretching vibrations, respectively).

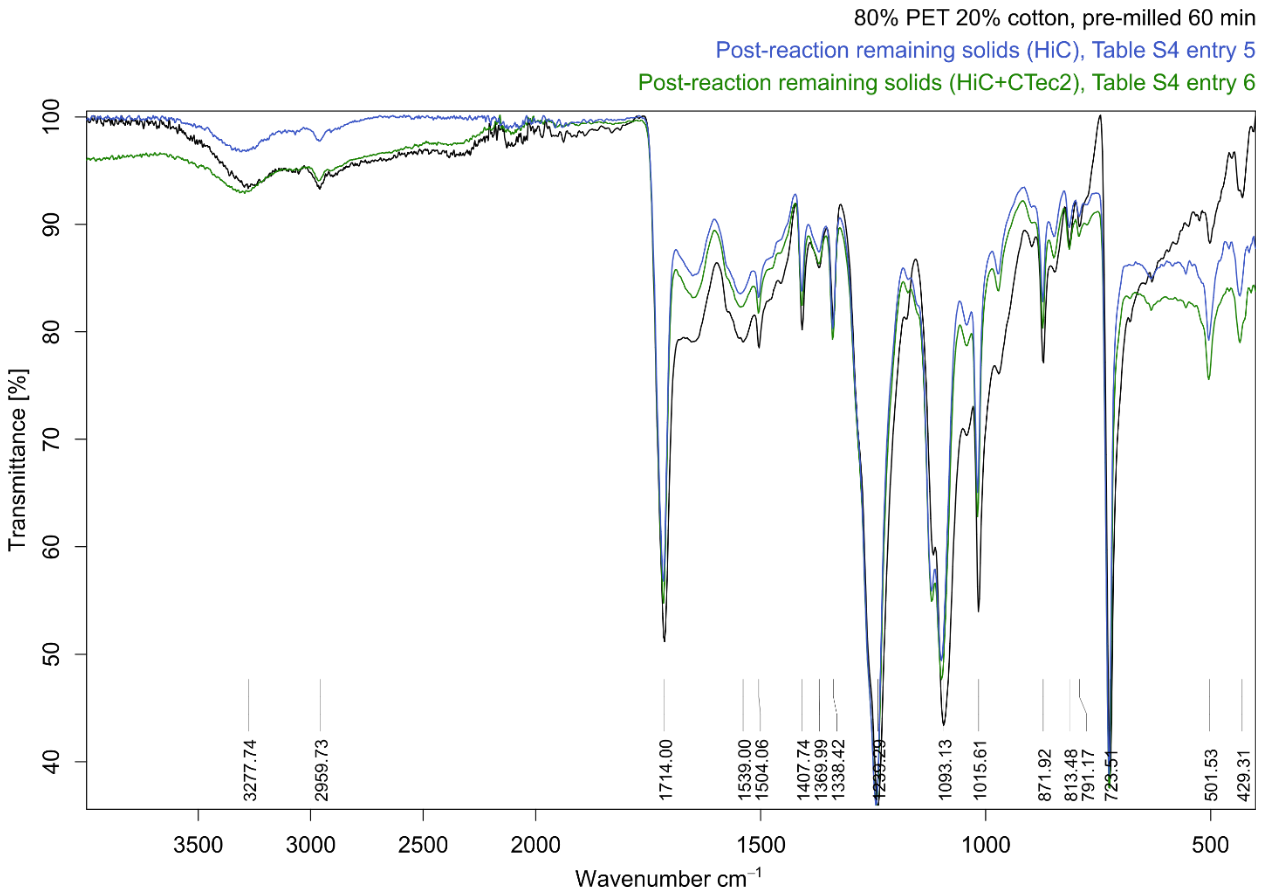

***Figure S9.***

FTIR spectra of the **PRE 80/20** pre-consumer white 80% PET and 20% cotton textile before (pre-milled textile, spectra shown in black) and after the mechanoenzymatic reaction (blue and green), shown in full recorded range. The post-hydrolysis washed materials show the difference of applying HiC enzyme alone (blue) or together with CTec2 cellulases (green), which lead to 14.1 ± 0.4% and 17 ± 2% weight loss, respectively.

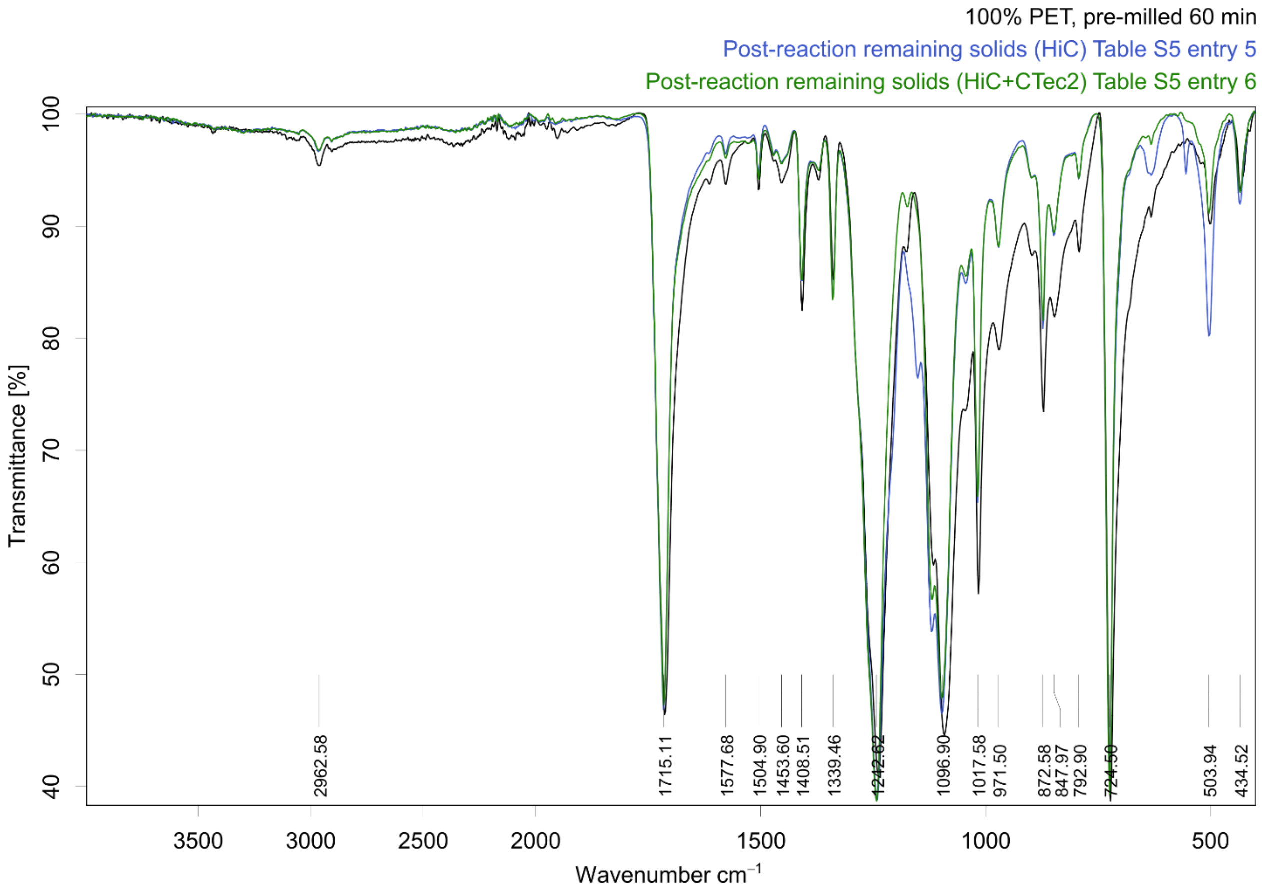

***Figure S10.***

FTIR spectra of the **PRE 100** pre-consumer white 100% PET textile before (pre-milled textile, spectra shown in black) and after the mechanoenzymatic reaction (blue and green), shown in full recorded range. The post-hydrolysis washed materials show the difference of applying HiC enzyme alone (blue) or together with CTec2 cellulases (green), which lead to equal 14 ± 3% and 14 ± 2% weight loss, respectively.

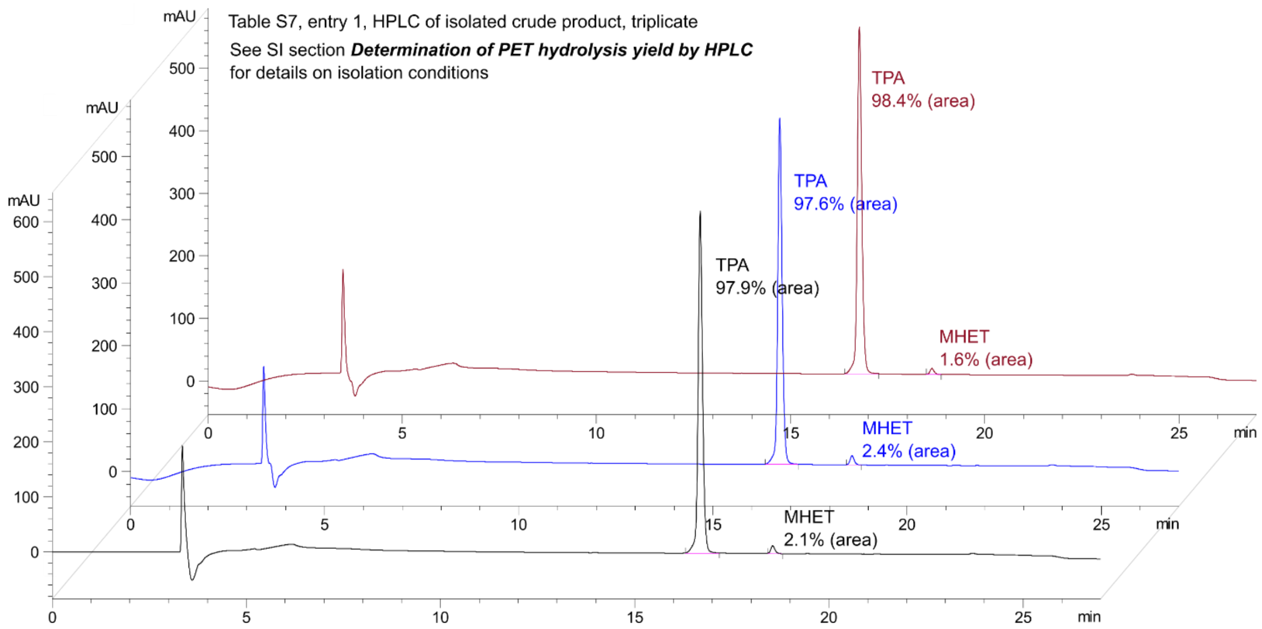

***Figure S11.***

HPLC chromatograms of the crude reaction products from multi-round optimized reactions, at conditions shown in Table S7, entry 1 (showing one triplicate). TPA selectivity over MHET is very high (50-fold), which considers the slightly higher extinction coefficient of MHET at the measurement condition (1.03x that of TPA), estimated based on the TPA and MHET calibration curves.

### Supplementary Tables

#### *Table S1.*

The varied conditions and results for the milling + aging reactions. Same for all reactions: the textile substrate was the pre-consumer white-colored 65% PET – 35% cotton textile **PRE 65/35**, the liquid-to-solid ratio was 1.5 µL $mg^{-1}$, and the buffer was 0.1 M NaPi pH 7.3.

| Entry | Textile (mg) | Pre-milling time | Enz loading HiC [i] | Enz loading CTec2 [ii] | Milling (min) | Aging (d) | **Yield of TPA (%)** | **Yield of glucose (%)** [iii] |
|---|---|---|---|---|---|---|---|---|
| 1 | 200 | None | 0.65 wt% (1 wt% to PET) | - | 5 | 7 | **1.8 ± 0.1** | **N/A** |
| 2 | 200 | None | 0.65 wt% (1 wt% to PET) | - | 30 | 7 | **12 ± 1** | **N/A** |
| 3 | 200 | None | 0.65 wt% (1 wt% to PET) | 0.7 wt% (2 wt% to cotton) | 5 | 7 | **1.4 ± 0.1** | **56.4 ± 0.6** |
| 4 | 200 | None | 0.65 wt% (1 wt% to PET) | 0.7 wt% (2 wt% to cotton) | 30 | 7 | **14 ± 1** | **61.2 ± 0.5** |
| 5 | 200 | None | - | 0.7 wt% (2 wt% to cotton) | 5 | 7 | **N/A** | **79 ± 2** |
| 6 | 200 | None | - | 0.7 wt% (2 wt% to cotton) | 30 | 7 | **N/A** | **81 ± 4** |
| 7 | 200 | 20 min | 0.65 wt% (1 wt% to PET) | 0.7 wt% (2 wt% to cotton) | 5 | 7 | **15 ± 1** | **52 ± 2** |
| 8 | 200 | 40 min | 0.65 wt% (1 wt% to PET) | 0.7 wt% (2 wt% to cotton) | 5 | 7 | **14.3 ± 0.7** | **56.1 ± 0.5** |
| 9 | 200 | 60 min | 0.65 wt% (1 wt% to PET) | 0.7 wt% (2 wt% to cotton) | 5 | 7 | **10.8 ± 0.6** | **58.4 ± 0.6** |

[i] Corresponds to 200 µL of the commercial HiC preparation (6.5 mg $mL^{-1}$ protein content) [ii] Corresponds to 20 µL of the CTec2 enzyme preparation (6.1 mg $mL^{-1}$ protein content). [iii] The amount of glucose added with the enzyme preparation (see ***Enzyme inherent glucose content determination***) was subtracted from the experimentally determined total glucose before calculating this yield.

***Table S2.***

The varied conditions and results for the milling + aging reactions on colored post-consumer **POST 65/35** and **POST 100** textiles. Same for all reactions: the liquid-to-solid ratio was 1.5 μL mg$^{-1}$ and the buffer added was 0.1 M NaPi pH 7.3.

| Entry | Textile (mg) | Textile composition | Pre-milling time | Enz loading HiC [i] | Enz loading CTec2 [ii] | Milling (min) | Aging (days) | **Yield of TPA (%)** | **Yield of glucose (%)** [iii] |
|---|---|---|---|---|---|---|---|---|---|
| 1 | 300 | 65% PET<br>35% cotton | None | 0.65 wt%<br>(1 wt% to PET) | - | **5** | 7 | **2.41 ± 0.03** | **N/A** |
| 2[iv] | 200 | 65% PET<br>35% cotton | None | 0.65 wt%<br>(1 wt% to PET) | 0.7 wt%<br>(2 wt% to cotton) | **5** | 7 | **1.38 ± 0.09** | **ND** |
| 3 | 300 | 65% PET<br>35% cotton | None | 0.65 wt%<br>(1 wt% to PET) | 0.7 wt%<br>(2 wt% to cotton) | **30** | 7 | **5.8 ± 0.5** | **41 ± 3** [v] |
| 4[iv] | 200 | 65% PET<br>35% cotton | **30 min** | 0.65 wt%<br>(1 wt% to PET) | 0.7 wt%<br>(2 wt% to cotton) | 5 | 7 | **8.2 ± 0.2** | **ND** |
| 5[iv] | 200 | 100% PET | None | 0.65 wt% | - | **5** | 7 | **3 ± 1** | **N/A** |
| 6 | 300 | 100% PET | None | 0.65 wt% | - | **30** | 7 | **7 ± 1** | **N/A** |
| 7[iv] | 200 | 100% PET | **30 min** | 0.65 wt% | - | 5 | 7 | **8.8 ± 0.2** | **N/A** |

[i] Corresponds to 200 μL of the commercial HiC preparation (6.5 mg mL$^{-1}$ protein content) [ii] Corresponds to 20 μL of the CTec2 enzyme preparation (6.1 mg mL$^{-1}$ protein content). [iii] The amount of glucose added with the enzyme preparation (see ***Enzyme inherent glucose content determination***) was subtracted from the experimentally determined total glucose before calculating this yield. [iv] Reaction carried out in duplicate. [v] The second batch of CTec2 was used in these reactions (see section ***Characterization of the enzymes***) and therefore the lower glucose yield of was affected by the lower CTec2 activity of this batch.

***Table S3.***

The varied conditions and results for Reactive Aging (RAging) reactions. Same for all reactions: the textile substrate was the pre-consumer white-colored 65% PET – 35% cotton textile **PRE 65/35** and the buffer added was 0.1 M NaPi pH 7.3.

| Entry | Textile (mg) | Pre-milling time | Enz loading HiC [i] | Enz loading CTec2 [ii] | η (μL mg$^{-1}$) | RAging regime (min) | No. of cycles | **Yield of TPA (%)** | **Yield of glucose (%)** [iii] | **Weight loss (%)** |
|---|---|---|---|---|---|---|---|---|---|---|
| 1[iv] | 200 | **20 min** | 0.65 wt% (1 wt% to PET) | - | 1.5 | 5 min + 1 day 55°C | 6 | **16.3 ± 0.6** | **ND** | **13.2 ± 0.4** |
| 2[iv] | 200 | **20 min** | 0.65 wt% (1 wt% to PET) | 0.7 wt% (2 wt% to cotton) | 1.5 | 5 min + 1 day 55°C | 6 | **12 ± 1** | **58 ± 3** | **36.6 ± 0.6** |
| 3 | 200 | **40 min** | 0.65 wt% (1 wt% to PET) | - | 1.5 | 5 min + 1 day 55°C | 6 | **15 ± 1** | **N/A** | **11.0 ± 0.9** |
| 4[v] | 200 | **40 min** | 0.65 wt% (1 wt% to PET) | 0.7 wt% (2 wt% to cotton) | 1.5 | 5 min + 1 day 55°C | 6 | **16 ± 3** | **50 ± 6** | **38 ± 2** |
| 5 | 200 | **40 min** | - | 0.7 wt% (2 wt% to cotton) | 1.5 | 5 min + 1 day 55°C | 1<br>3<br>6 | **N/A** | **53 ± 2**<br>**71 ± 6**<br>**63 ± 6** | **ND**<br>**ND**<br>**32 ± 2** |
| 6 | 200 | **60 min** | 0.65 wt% (1 wt% to PET) | - | 1.5 | 5 min + 1 day 55°C | 6 | **16.7 ± 0.6** | **N/A** | **11.0 ± 0.9** |
| 7 | 200 | **60 min** | 0.65 wt% (1 wt% to PET) | 0.7 wt% (2 wt% to cotton) | 1.5 | 5 min + 1 day 55°C | 6 | **17 ± 2** | **56 ± 3** | **39 ± 2** |
| 8 | 200 | 40 min | **dialyzed** [vi] 0.65 wt% (1 wt% to PET) | 0.7 wt% (2 wt% to cotton) | 1.5 | 5 min + 1 day 55°C | 6 | **10 ± 1** | **70 ± 5** | **39 ± 3** |
| 9 | 200 | 40 min | 0.65 wt% (1 wt% to PET) | 0.7 wt% (2 wt% to cotton) | **2** | 5 min + 1 day 55°C | 6 | **16 ± 3** | **54 ± 3** | **33 ± 2** |
| 10 | 200 | 40 min | 0.65 wt% (1 wt% to PET) | 0.7 wt% (2 wt% to cotton) | **2.5** | 5 min + 1 day 55°C | 6 | **13 ± 1** | **61 ± 1** | **34 ± 1** |
| 11 | 200 | 40 min | 0.65 wt% (1 wt% to PET) | 0.7 wt% (2 wt% to cotton) | **3** | 5 min + 1 day 55°C | 6 | **15 ± 2** | **77 ± 10** | **32 ± 5** |
| 12 | 200 | 40 min | - | 0.7 wt% (2 wt% to cotton) | 1.5 | **5 min + 1 h 55°C** | 1<br>3<br>6<br>10 | **N/A** | **10.4 ± 0.5**<br>**17.1 ± 0.1**<br>**19.7 ± 0.5**<br>**22 ± 2** | **ND** |

[i] Corresponds to 200 μL of the commercial HiC preparation (6.5 mg mL$^{-1}$ protein content) [ii] Corresponds to 20 μL of the CTec2 enzyme preparation (6.1 mg mL$^{-1}$ protein content). [iii] The amount of glucose added with the enzyme preparation (see ***Enzyme inherent glucose content determination***) was subtracted from the experimentally determined total glucose before calculating this yield. [iv] Reaction performed in duplicate. [v] reaction performed in quintuplicate. [vi] The dialyzed HiC powder was prepared as described in ***Dialysis and lyophilization of the commercial Novozym® 51032 cutinase solution***.

***Table S4.***

The varied conditions and results for Reactive Aging (RAging) reactions. Same for all reactions: the textile substrate here was the pre-consumer white-colored 80% PET 20% cotton textile **PRE 80/20**, the liquid-to-solid ratio was 1.5 µL mg$^{-1}$ and the buffer added was 0.1 M NaPi pH 7.3.

| Entry | Textile (mg) | Pre-milling time | Enz loading HiC [i] | Enz loading CTec2 [ii] | RAging regime (min) | Number of cycles | **Yield of TPA (%)** | **Yield of glucose (%)** [iii] | **Weight loss (%)** |
|---|---|---|---|---|---|---|---|---|---|
| 1[iv] | 200 | 20 min | 0.65 wt% (0.8 wt% to PET) | - | 5 min + 1 day 55°C | 6 | **8.3 ± 0.9** | **N/A** | **15.4 ± 0.8** |
| 2[iv] | 200 | 20 min | 0.65 wt% (0.8 wt% to PET) | 0.7 wt% (3.5 wt% to cotton) | 5 min + 1 day 55°C | 6 | **9.5 ± 0.6** | **0** | **17.6 ± 0.8** |
| 3 | 200 | 40 min | 0.65 wt% (0.8 wt% to PET) | - | 5 min + 1 day 55°C | 6 | **7.0 ± 0.1** | **N/A** | **15 ± 2** |
| 4 | 200 | 40 min | 0.65 wt% (0.8 wt% to PET) | 0.7 wt% (3.5 wt% to cotton) | 5 min + 1 day 55°C | 6 | **7.9 ± 0.6** | **0** | **17 ± 3** |
| 5 | 200 | 60 min | 0.65 wt% (0.8 wt% to PET) | - | 5 min + 1 day 55°C | 6 | **7.8 ± 0.8** | **N/A** | **14.1 ± 0.4** |
| 6 | 200 | 60 min | 0.65 wt% (0.8 wt% to PET) | 0.7 wt% (3.5 wt% to cotton) | 5 min + 1 day 55°C | 6 | **9.2 ± 0.6** | **0** | **17 ± 2** |

[i] Corresponds to 200 µL of the commercial HiC preparation (6.5 mg mL$^{-1}$ protein content) [ii] Corresponds to 20 µL of the CTec2 enzyme preparation (6.1 mg mL$^{-1}$ protein content). [iii] The amount of glucose added with the enzyme preparation (see ***Enzyme inherent glucose content determination***) was subtracted from the experimentally determined total glucose before calculating this yield. [iv] Reaction performed in duplicate.

***Table S5.***

The varied conditions and results for Reactive Aging (RAging) reactions. Same for all reactions: the textile substrate here was the pre-consumer white-colored 100% PET textile **PRE 100**, the liquid-to-solid ratio was 1.5 μL $mg^{-1}$ and the buffer added was 0.1 M NaPi pH 7.3.

| Entry | Textile (mg) | Pre-milling time | Enz loading HiC [i] | Enz loading CTec2 [ii] | RAging regime (min) | Number of cycles | **Yield of TPA (%)** | **Weight loss (%)** |
|---|---|---|---|---|---|---|---|---|
| 1[iii] | 200 | 20 min | 0.65 wt% | - | 5 min + 1 day 55°C | 6 | **11 ± 2** | **15 ± 1** |
| 2[iii] | 200 | 20 min | 0.65 wt% | 0.7 wt% | 5 min + 1 day 55°C | 6 | **11 ± 1** | **14.9 ± 0.7** |
| 3 | 200 | 40 min | 0.65 wt% | - | 5 min + 1 day 55°C | 6 | **11 ± 1** | **15 ± 1** |
| 4[iii] | 200 | 40 min | 0.65 wt% | 0.7 wt% | 5 min + 1 day 55°C | 6 | **11.6 ± 0.7** | **16.3 ± 0.4** |
| 5 | 200 | 60 min | 0.65 wt% | - | 5 min + 1 day 55°C | 6 | **11 ± 1** | **14 ± 3** |
| 6 | 200 | 60 min | 0.65 wt% | 0.7 wt% | 5 min + 1 day 55°C | 6 | **12 ± 1** | **14 ± 2** |

[i] Corresponds to 200 μL of the commercial HiC preparation (6.5 mg $mL^{-1}$ protein content) [ii] Corresponds to 20 μL of the CTec2 enzyme preparation (6.1 mg $mL^{-1}$ protein content). [iii] Reaction performed in duplicate.

***Table S6.***

The varied conditions and results for Reactive Aging (RAging) reactions. Same for all reactions: the textile substrate was the colored post-consumer 65% PET - 35% cotton textile **POST 65/35**, the liquid-to-solid ratio was 1.5 μL $mg^{-1}$ and the buffer added was 0.1 M NaPi pH 7.3.

| Entry | Textile (mg) | Pre-milling time | Enz loading HiC [i] | Enz loading CTec2 [ii] | RAging regime (min) | Number of cycles | **Yield of TPA (%)** | **Yield of glucose (%)** [iii] | **Weight loss (%)** |
|---|---|---|---|---|---|---|---|---|---|
| 1[iv] | 200 | 30 min | 0.65 wt% (1 wt% to PET) | - | 5 min + 1 day 55°C | 6 | **9 ± 1** | **N/A** | **10.4 ± 0.5** |
| 2[iv] | 200 | 30 min | 0.65 wt% (1 wt% to PET) | 0.7 wt% (2 wt% to cotton) | 5 min + 1 day 55°C | 6 | **11 ± 1** | **33 ± 4** [v] | **31 ± 7** |

[i] Corresponds to 200 μL of the commercial HiC preparation (6.5 mg $mL^{-1}$ protein content) [ii] Corresponds to 20 μL of the CTec2 enzyme preparation (6.1 mg $mL^{-1}$ protein content). [iii] The amount of glucose added with the enzyme preparation (see ***Enzyme inherent glucose content determination***) was subtracted from the experimentally determined total glucose before calculating this yield. [iv] Reaction was performed in a duplicate of triplicates, by two members of the research lab. The TPA yield and weight loss, with the corresponding standard deviations, are calculated based on all 6 reactions per entry, but the glucose yield was measured only for one triplicate of entry 2 and is therefore calculated based on 3 reactions. [v] The second batch of CTec2 was used in these reactions (see section ***Characterization of the enzymes***) and therefore the lower glucose yield of was affected by the lower CTec2 activity of this batch.

***Table S7.***

Blank (no enzyme) control reactions for the pre-milled textiles using the Reactive Aging (RAging). Same for all reactions: all textiles were pre-milled 20–60 min, the liquid-to-solid ratio was 1.5 $\mu L\ mg^{-1}$ and the buffer added was 0.1 M NaPi pH 7.3, no enzymes were added.

| Entry | Textile (mg) | Textile | RAging regime (min) | Number of cycles | **Yield of TPA (%)** | **Yield of glucose (%)** | **Weight loss (%)[ii]** |
|---|---|---|---|---|---|---|---|
| 1[i] | 200 | **PRE 65/35** | 5 min + 1 day 55°C | 6 | **0** | **ND** | **0** |
| 2[i] | 200 | **PRE 80/20** | 5 min + 1 day 55°C | 6 | **0.08** | **ND** | **0** |
| 3[i] | 200 | **PRE 100** | 5 min + 1 day 55°C | 6 | **0.07** | **ND** | **0** |
| 4[i] | 200 | **POST 65/35** | 5 min + 1 day 55°C | 6 | **0** | **ND** | **0** |

[i] Reaction performed in monoplicate. [ii] Weight loss was determined as described in section ***Determining the weight loss of textiles after mechanoenzymatic hydrolytic reactions***, with the exception that the dry weight of each reaction mixture on day 6 was calculated based on the solid loading (40 wt%), an not based on the initial drying step. Washing steps were performed identically to reactions, where enzymes were employed.

***Table S8.***

The varied conditions and results for multi-round Reactive Aging (RAging) reactions (see section ***Method for the multi-round RAging reactions with fresh enzyme additions***). Same for all reactions: the textile was pre-consumer white-colored 65% PET – 35% cotton textile **PRE 65/35**, pre-milled 40 minutes at 30 Hz, the liquid-to-solid ratio was kept constant at 1.5 μL $mg^{-1}$ and the buffer was 0.1 M NaPi pH 7.3.

| | | 1st round | | | | | 2nd round – 5th round | | | | | Total 1st – 5th round |
|---|---|---|---|---|---|---|---|---|---|---|---|---|
| Entry | Textile (mg) | Enz loading | **RAging regime (1st round)** | **Yield of glucose (%)** | Washed after round? | Weight loss (%) | Enz loading | RAging regime (1 round) | **Round no.** | **Yield of TPA (%)** | Washed after each round? | Weight loss (%) |
| 1 | 600 | CTec2 [i] 0.7 wt% (2 wt% to cotton) | 3 cycles: 5 min + 1 day 55°C | **83 ± 4** [ii] | no | ND | dialyzed HiC [iii] 0.65 wt% | 5 cycles: 5 min + 1 day 55°C | **2.**<br>**3.**<br>**4.**<br>**5.** | **9.0 ± 0.8**<br>**14.6 ± 0.8**<br>**21 ± 2**<br>**28 ± 2** | no | ND |
| 2 | 600 | CTec2 [i] 0.7 wt% (2 wt% to cotton) | 3 cycles: 5 min + 1 day 55°C | **68.6 ± 0.2** [ii] | yes, milliQ | 31 ± 1 | HiC [iv] 0.65 wt% | 5 cycles: 5 min + 1 day 55°C | **2.**<br>**3.**<br>**4.**<br>**5.** | **11.4 ± 0.9**<br>**19 ± 1**<br>**26 ± 2**<br>**30 ± 2** | yes, sodium carbonate, milliQ | 62.4 ± 0.2 |

[i] Corresponds to 60 μL of the CTec2 enzyme preparation (6.1 mg $mL^{-1}$ protein content). [ii] The amount of glucose added with the enzyme preparation (see ***Enzyme inherent glucose content determination***) was subtracted from the experimentally determined total glucose before calculating this yield. [iii] The lyophilized HiC was prepared as described in ***Dialysis and lyophilization of the commercial Novozym® 51032 cutinase solution***. [iv] The amount of the added commercial HiC preparation (6.5 mg $mL^{-1}$ protein content) was chosen based on the weight of the remaining material after washing and drying, at the start of each cycle (rounds 2–5).
